# Nicotinamide combined with gemcitabine is an immunomodulatory therapy that restrains pancreatic cancer in mice

**DOI:** 10.1101/2020.06.02.129809

**Authors:** Benson Chellakkan Selvanesan, Kiran Meena, Amanda Beck, Lydie Meheus, Olaya Lara, Ilse Rooman, Claudia Gravekamp

## Abstract

Treatments for pancreatic ductal adenocarcinoma (PDAC) are poorly effective, at least partly due to the tumor’s immune-suppressive stromal compartment. New evidence of positive effects on immune responses in the tumor microenvironment, compelled us to test the combination of gemcitabine (GEM), a standard chemotherapeutic for pancreatic cancer, with nicotinamide (NAM), the amide form of niacin (vitamin B3), in various mouse tumor models of pancreatic cancer, i.e. peritoneal or orthotopic of Panc-02 (Kras^G12D^) and orthotopic KPC (Kras^G12D^, p53^R172H^, Pdx1-Cre) grafts. A significant reduction in tumor weight and number of metastases was found, as well as a significant improved survival of the NAM+GEM group compared to all control groups. Immunohistochemistry and flow cytometry of pancreatic tumors showed a significant decrease in tumor-associated macrophages (TAM) and myeloid-derived suppressor cells (MDSC), an increase in the number of CD4 and CD8 T cells and the production of granzyme B in the NAM+GEM group. Moreover, T cell responses to tumor-associated antigen survivin were observed in spleens of the mice that received NAM+GEM but not in those that received single agents or saline. In addition, remodeling of the tumor stroma was observed with decreased collagen I and expression of hyaluronic acid binding protein, reorganization of the immune cells into lymph node like structures, and CD31 positive vessels. Expression profiling for a panel of immuno-oncology genes revealed significant changes in genes involved in migration and activation of T cells, attraction of dendritic cells, and epitope spreading. This study highlights the potential of NAM+GEM as immunotherapy for advanced pancreatic cancer.

## INTRODUCTION

Patients with pancreatic ductal adenocarcinoma (PDAC) have a five-year survival below 10%. Part of this poor outcomes has been attributed to the fact that PDAC is characterized by a dense desmoplastic stroma that prevents drugs and immune cells from penetrating the pancreatic tumors^1,2^. This stroma consists of different types of cancer activated fibroblasts (CAF)^3^ that not only produce extracellular matrix (ECM) components such as collagen but also crosstalk to immune cells^4,5^. Among these inflammatory cells are tumor-associated macrophages (TAM) and myeloid-derived suppressor cells (MDSC), that promote immune suppression, tumor progression, angiogenesis, invasion, and extravasation of tumor cells resulting in the development of metastases^6^. Immunomodulatory therapies to counteract tumor progression are highly sought after but with limited success.

Gemcitabine (GEM) is an FDA-approved drug for advanced pancreatic cancer (PDAC) that in combination with nab-paclitaxel^7,8^, only modestly improves patient survival^9–11^. Hence, more effective therapies are warranted. GEM is a deoxycytidine analogue that inhibits DNA synthesis. New evidence indicates that antitumor activities of chemotherapy such as GEM also rely on several off-target effects, especially directed to the host’s immune system that contribute to tumor eradication. So far, studies that have been focusing on evaluating the effect of GEM on T cell responses to the tumors are limited. A few reports describe that GEM improves T cell activation in mice and humans with pancreatic cancer^12–16^. GEM is known for its ability to reduce the MDSC population in tumor-bearing humans and mice^17^. Finding drugs to combine with GEM that can strengthen the immune response would be a significant step forward.

Nicotinamide (NAM), the amide form of niacin (vitamin B3), is a nutrient provided by dietary source and supplement, and has little side effects^18^. NAM is a precursor of the coenzyme nicotinamide adenine dinucleotide (NAD^+^) and participates in the cellular energy metabolism in the mitochondrial electron transport chain^19^. It has been shown that NAM kills pancreatic tumor cells through down regulation of SIRT-1, K-ras, and Akt-1 expression^20^, and that NAM sensitizes tumor cells to chemo- and radiotherapy through inhibition of poly (ADP-ribose) polymerase (PARP)^21^. New evidence has been reported that NAM affects immune responses positively. For instance, NAM increases the expansion of human T cells through mitochondrial activation^22^. Also, peritumoral and infiltrating CD4 and CD8 T cells were significantly increased in melanomas upon NAM treatment compared to a placebo^23^. Together this suggests that NAM may qualify as an effective therapeutic add-on in PDAC. Moreover, it is safe to use and showed cancer preventative actions in a phase 3 clinical trial of non-melanoma skin cancer ^24^.

Based on this information we evaluated the effect of combining GEM with NAM on pancreatic cancer and focused on the immune system. We found a significant decrease in the growth of primary tumors and metastases, as well as an increase in the survival time of NAM+GEM treated mice, in correlation with a significant influx of immune cells into the pancreatic tumors, and with a significant increase in CD31-positive blood vessels. The immune infiltrate was characterized by T cells that formed peritumoral lymph node-like structures (LNS) predominantly in the NAM+GEM group, and intratumoral LNS sometimes in the NAM or GEM groups. CD4 and CD8 T cells were activated by NAM+GEM to the tumor-associated antigen (TAA) survivin, which is highly expressed by tumors and metastases in the Panc-02 model^25^, one of the experimental models used here. In addition, the number of TAM and MDSC in the tumor decreased. Potential immunoregulatory mechanisms of NAM+GEM based on Nanostring analysis will be discussed.

## MATERIAL AND METHODS

### Animals

C57BL/6 female mice aged 3 months were obtained from Charles River. These mice were used to generate the Panc-02 model. Male and female KPC-wild type mice were maintained at Einstein, which have the same genetic background as KPC mice^26^ but are negative for Kras^G12D^ and Tp53^R172H^ mutations and for Cre-recombinase. These mice were used to generate the orthotopic KPC mice.

### Cell lines

The Panc-02 cell line with the Kras^G12D^ mutant allele was derived from a methylcholanthrene-induced ductal adenocarcinoma growing in a C57BL/6 female mouse^27^. Panc-02 cells were cultured in McCoy’s5A medium supplemented with 10% fetal bovine serum, 2 mM glutamine, non-essential amino acids, 1 mM sodium pyruvate, and 100 U/ml Pen/Strep. The KPC tumor cell line was derived from a pancreatic KPC tumor (*LSL^KrasG12D+/^ LSL-p53^R172-/-^; Pdx1-Cre*) in our lab, which was kindly by Jacco van Rheenen, Cancer Genomics, Utrecht, The Netherlands^28^.

### Tumor development

Three different mouse tumor models have been used, i.e. a peritoneal Panc-02 model, an orthotopic Panc-02 model, and an orthotopic KPC model. **(Fig S1).**

### Peritoneal Panc-02 model

Panc-02 tumor cells were injected into the mammary fat pad (peritoneal cavity model) (10^5^) of C57BL/6 mice. Ten days later a small primary tumor has grown into peritoneal membrane (average tumor weight is 0.03 grams which equals ≈5 mm^2^)**(Fig S2),** and tumor cells metastasize via the peritoneal cavity to other organs (average of 50 small metastases visible by the naked eye), predominantly to the pancreas and liver, and less abundantly to mesenchymal lymph nodes (LN) along the gastro-intestines (GI) and the diaphragm^29^. This model has been named peritoneal Panc-02 because metastases migrate directly from the primary tumor via the peritoneal cavity to the pancreas, liver and mesenteric lymph nodes along the GI. In the pancreas the metastases cluster together. The tumor weight was measured and the number of metastases was counted after euthanizing the mice. This model has significantly more metastases than the orthotopic models, particularly in the liver, and therefore produce more ascites than the orthotopic models.

### Orthotopic Panc-02 model

Orthotopic Panc-02 tumors were generated in C57BL/6 mice as we described previously^16^. Briefly, mice were anesthetized with ketamineb (Mylan Institutional LLC)/xylazine (Akorn Animal Health)(respectively 100 mg and 10 mg/kg, ip) the hair was removed at the location of the spleen, and the skin was sterilized with betadine followed by 70% alcohol. The animal was covered with gauze sponge surrounding the incision site. A 1 cm incision was made in the abdominal skin and muscle just lateral to the midline and directly above the spleen/pancreas to allow visualization. The spleen/pancreas was gently retracted and positioned to allow injection of Pan-02 tumor cells (10^6^/50μL PBS) directly into the pancreas, from the tail all the way to the head of the pancreas. To prevent leakage of injected cell suspension, the injection site was tied off after tumor cell injections with dissolvable suture. The spleen/pancreas were then replaced within the abdominal cavity, and both muscle and skin layers closed with sutures. Following recovery from surgery, mice were monitored and weighed daily. A palpable tumor appeared within 10 days in the pancreas. In this model, tumor cells migrate via the blood stream to the liver and grow into small metastases visible by the naked eye. Sometimes, some tumor cells leaked into the peritoneal cavity. Matrigel was not used here since it may prevent dissemination of tumor cells and the development of metastases. The tumor weight was measured and the number of metastases was counted after euthanizing the mice.

### Orthotopic KPC model

Orthotopic KPC tumors were generated similarly, but now the KPC tumor cells (10^5^/50μL PBS) were injected into the pancreas of “KPC-negative” mice. KPC-negative mice have the same genetic background as the KPC transgenic mice (C57BL/6xFVBxJ129) but lack the expression of Kras and p53 mutations and the Pdx-Cre, and allow the generation of pancreatic tumors and metastases in the liver. In this orthotopic model, a palpable tumor appeared within 10 days, and the metastases are blood born, like in the orthotopic Panc-02 model.

### Protocol for treatment of mice with NAM+GEM

The optimal dose of NAM (Green Labs Nutrition, Poland) was determined by testing different concentrations of NAM (100, 50 and 25, 12.5 and 6.25 mg/200 μL/dose). The highest dose without any physiological side effects appeared to be 25 mg/200 μL. Based on these results, we decided to use 20 mg/200μL as the optimal dose used in all experiments. First tumors and metastases were generated as described above. NAM (20/mg/200 μL/dose) was administered orally and GEM (1.2 mg/200 μL/dose) was administered ip, alternately, starting 10 days after tumor cell injection and continued for 2 weeks as outlined for all three pancreatic cancer models in **Fig S1abc**. Preparation of NAM: 1 gram of NAM was dissolved in 10 mls of apple juice. 16 mg of black pepper was added to the solution to prevent glucuronidation. One dose consisted of 20 mg of NAM in 200 μL apple juice. The NAM solution was aliquoted and stored at −20°C until use. Preparation of GEM (Fresensius Kabi; Gemita): 1 gram of GEM was dissolved in 100 mls of endotoxin-free saline, diluted until 6 mg/ml, and then aliquoted and stored at −20°C until use. One dose consisted of 1.2 mg in 200 μL saline.

### Flow Cytometry

Immune cells from spleens of mice were isolated as described previously^30^. Immune cells were also isolated from pancreatic tumors using Dispase (Roche cat#049422078001) and Collagenase (Sigma-Aldrich cat #C0130) as we described previously^31^. Anti-CD3 (BD Bioscience, cat # 560527) and anti-CD8 antibodies (BD Bioscience, cat # 552877) were used to identify CD8 T cells, anti-CD3 and anti-CD4 (BD Bioscience, cat # 552051) to identify CD4 T cells, anti-CD11b (BD Bioscience, cat # 553312) and anti-Gr1 (BD Bioscience, cat # 553127) to identify MDSC, and anti-CD11b and anti-F4/80 (eBioscience, cat # 17-4801-82) to identify TAM. Appropriate isotype controls were included for each sample. 50,000-100,000 cells were acquired by scanning using a special order LSR-II Fluorescence Activated Cell Sorter system (Beckton and Dickinson), and analyzed using FlowJo 7.6 software. Cell debris and dead cells were excluded from the analysis based on scatter signals and use of Fixable Blue or Green Live/Dead Cell Stain Kit (BD Bioscience, cat # 564406).

### ELISPOT

Immune cells from spleens or tumors were isolated from treated and control Panc-02 mice for ELISPOT (BD Biosciences, cat# 551083) analysis, as described previously^31,32^. To detect T cell responses to survivin, 10^5^ spleen cells were transfected with pcDNA-3.1-survivin or pcdNA3.1 alone (negative control) or nothing (negative control). The frequency of IFNγ-producing spleen cells was measured 72 hrs later using an ELISPOT reader (CTL Immunospot S4 analyzer). To determine the frequency of IFNγ-producing CD4 and CD8 T cells, spleen cells were depleted for CD4 and CD8 T cells using magnetic bead depletion techniques according to the manufacturer’s instructions (Miltenyi Biotec).

### Immunohistochemistry (IHC)

Tumors were dissected from pancreas and immediately fixed with buffered formalin, and the tissue was embedded in paraffin. Sections (5 μm) were sliced and placed on slides, then deparaffinized at 60°C for 1 hr, followed by xylene, an ethanol gradient (100-70%), water, and PBS. Slides were then incubated for 30 min in 3% hydrogen peroxide followed by boiling in citrate buffer for 20 min. Once the slides were cooled, washed, and blocked with 5% goat serum, the sections were incubated with primary antibodies such as anti-CD4 (Cell Signaling Technology, cat # 25229) (1:100 dilution), anti-CD8α (1:400 dilution)(Cell Signaling Technology, cat # 98941), anti-Perforin (Cell Signaling Technology, cat # 31647)(1:300 dilution), anti-Granzyme (Cell Signaling Technology, cat # 44153) (1:200 dilution), anti-CD31 (Cell Signaling Technology, cat # 77699) (1:100 dilution), or anti-αSMA antibodies (Cell Signaling Technology, cat # 19245), followed by incubation with secondary antibody (mouse anti-goat IgG-HRP) (Cell Signaling Technology, cat# 8114S), and SignalStain^®^ Boost IHC Detection Reagent (Cell Signaling Technology, cat# 8114S). Subsequently, the slides were incubated with 3,3’-diaminobenzidine (DAB)(Vector Laboratories, cat# SK-4100), counterstained with hematoxylin, dehydrated through an ethanol gradient (70-100%) and xylene, and mounted with Permount. The slides were scanned with a 3D Histech P250 High Capacity Slide Scanner to acquire images and quantification data. Secondary antibodies without primary antibodies were used as negative control.

### Nanostring Technology

Pieces of 3 mm^3^ tumors were submerged in 5 vol of RNAlater (Invitrogen) (n = 5 samples/group). Total RNA was isolated from these tissues using TRIzol (Life Technologies) as we described previously^32^. The RNA concentrations in the samples were measured using a NanoDrop 2000 instrument (Thermo Fisher Scientific) and checked for quality on agarose gels. All RNA samples used in this study exhibited optical density (OD)260/280 ratios greater than 1.9 and RNA integrity numbers (RINs) greater than 8.5.

A total of 770 immune-related mouse genes were analyzed the nCounter Mouse PanCancer Immune Profiling Panel (NanoString) categorized in various pathways **(Table S1).**

### Bioinformatics Analysis

Differential expression analyses of mRNA expression data in 20 samples were performed by using the DESeq2 R package v1.20.0. A total of 770 immunology-related mouse genes, created from the nCounter Mouse PanCancer Immune Profiling Panel (NanoString), were then implemented as candidate genes in this study. GSVA was employed to detect the variation values of the GO term pathways in each group using the R package GSVA.

### Statistical Analysis Nanostring

All experiments were repeated in triplicate unless otherwise stated. Statistical analyses were performed using GraphPad Prism softwareversion 7 (GraphPad Software), and statistical significance was defined as p < 0.05. Two-way analysis of variance (ANOVA) was performed on the experimental data for tumor volumes and mouse weights. The results are presented as the mean ± standard error of the mean (SEM).

Differentially expressed genes were compared with Student**’**s t test, and adjusted p values of less than 0.01 and fold changes of greater than 2 were considered to indicate significant dysregulation. p values were adjusted by the Benjamini-Hochberg (BH) method. Adjusted p values are also called q values. Data were analyzed using R (version 3.5.1).

### Statistical Analysis tumors, metastases, and immunological responses

To statistically compare the effects NAM+GEM on the growth of metastases and tumors or on immune responses in the mouse models, the Mann-Whitney was applied using Prism. *p<0.05, **<0.01, ***<0.001, ****<0.0001. Values of p<0.05 were considered statistically significant. Survival studies were analyzed using Mantel-Cox test. p<0.05 is significant.

## RESULTS

### NAM+GEM significantly improves clinical parameters, including survival, in mouse PDAC

Here, we tested low doses of NAM (20 mg/dose) and GEM (1.2 mg/dose) in three different cancer models of pancreatic cancer, a peritoneal Panc-02 model, an orthotopic Panc-02 model, and an orthotopic KPC model **(Fig S1abc).** In the peritoneal Panc-02 model **(Fig S1a),** C57Bl6 mice were injected with 10^5^ Panc-02 tumor cells in the mammary fat pad, and 10 days later, when a tumor has grown into the peritoneal membrane and metastases developed predominantly in pancreas and liver **(Fig S2),** treatments were started and continued for 14 days as outlined in **Fig S1a** (4 treatments with NAM and 4 with GEM, alternately). At the end of treatment, mice were euthanized and analyzed for tumors and metastases. NAM+GEM reduced the tumor weight by 70% (p<0.01) and the number of metastases by 95% compared to the saline group (p<0.001) **(Fig 1ab).** Moreover, ascites production was observed in all groups (grade of ascites production 2-5; 5 is worst and 0 is no ascites) but was absent in the NAM+GEM group **(Fig 1c).** Pictures of metastases in the pancreas and liver in each treatment group are shown in **Fig S3ab.** We also compared the effect of NAM+GEM with anti-PD-L1 to NAM+GEM alone. As shown in **Fig 1ab,** anti-PD-L1+NAM+GEM was less effective against tumors and metastases than NAM+GEM alone. Why anti-PD-L1 had a negative on the NAM+GEM effect on the pancreatic cancer needs to be further analyzed.

**Figure 1:**
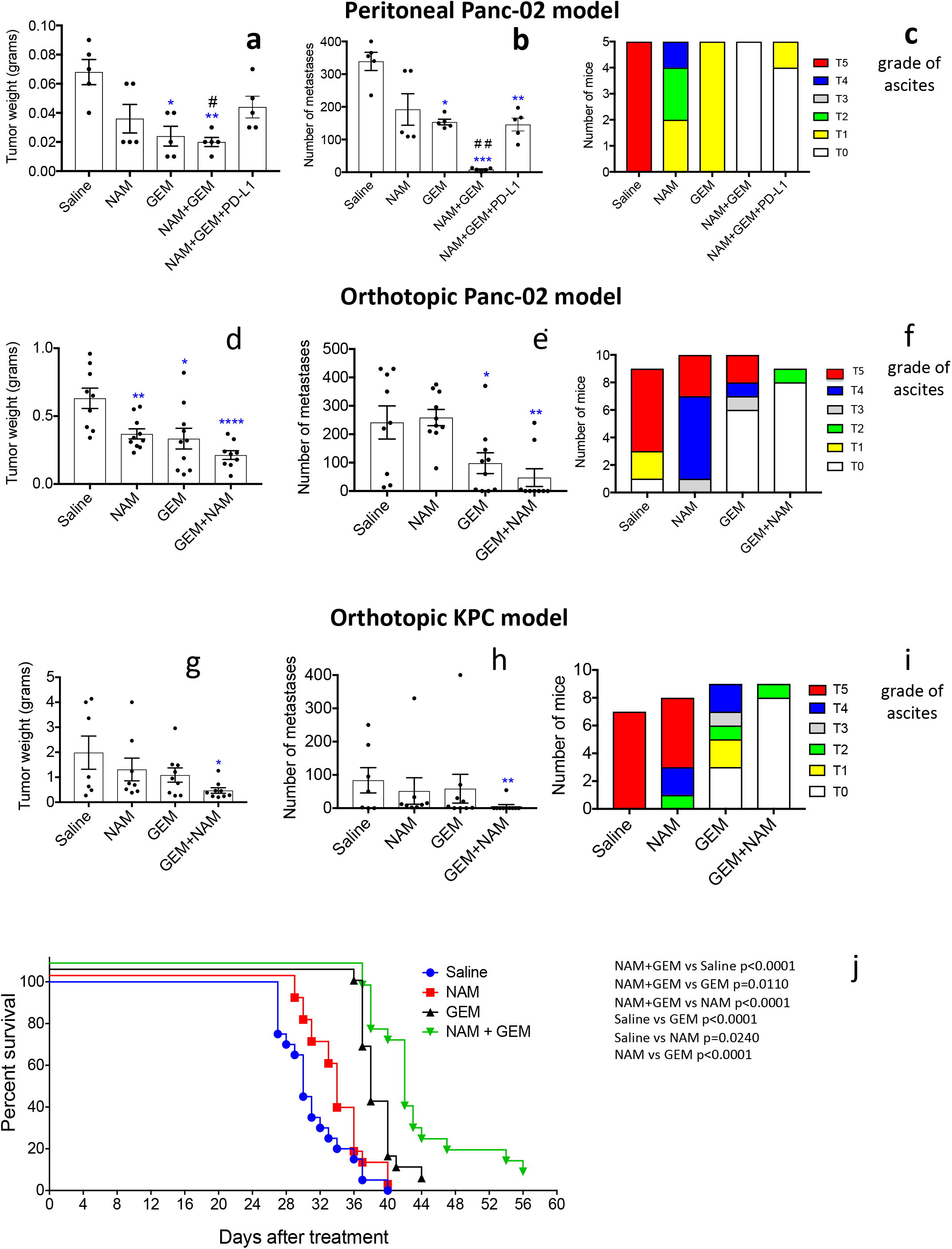
NAM+GEM significantly improves clinical parameters, including survival, in mouse PDAC. Peritoneal Panc-02 model. Tumors and metastases were generated and treatments with NAM+GEM and controls were performed as outlined in **Fig S1a.** At the conclusion of the experiment tumor weight **(a)** and number of metastases **(b)** was measured, as well as ascites **(c).** Representative of 2 experiments. Results were averaged with n=5 mice per group. For comparison, anti-PD-L1 antibody was added to the NAM+GEM therapy. **Orthotopic Panc-02 model.** Tumors and metastases were generated and treatments with NAM+GEM and controls were performed as outlined in **Fig S1b.** At the conclusion of the experiment tumor weight **(d)** and number of metastases **(e)** was measured, as well as ascites **(f).** Average of two experiments with n=10 mice per group. **Orthotopic KPC model.** Tumors and metastases were generated and treatments with NAM+GEM and controls were performed as outlined in **Fig S1c.** At the conclusion of the experiment tumor weight **(g)** and number of metastases **(h)** was measured, as well as ascites **(i).** Significant differences were determined by Mann-Whitney *p<0.05, **p<0.01, ***p<0.001, ****p<0.0001. Error bars represent standard error of the mean (SEM). **Survival study orthotopic Panc-02 model.** Tumors and metastases were generated and treatments with NAM+GEM and controls were performed as outlined in **Fig S1b.** At the end of treatments mice were monitored until death, and survival was assessed **(j)** when define clinical endpoints were reached (see M and M). The results of 2 experiments were averaged with n=10 mice per group. Significant differences were determined by the Mantel-Cox test. NAM=Nicotinamide, GEM=Gemcitabine.

Subsequently, we tested NAM+GEM in an orthotopic Panc-02 model. Briefly, 10^6^ tumor cells were orthotopically injected into the pancreas and 10 days later the treatment with NAM+GEM started as outlined in **Fig S1b** (at this time point tumors are palpable in the pancreas) and the effect of treatment on tumors and metastases was analyzed. As shown in **Fig 1de**, a significant reduction in tumor weight (p<0.0001) and metastases (p<0.01) was observed in the NAM+GEM group compared to the saline group, while ascites production was relatively high in all control groups (Grade 0-5) but relatively minor in the NAM+GEM group (Grade 0-2)**(Fig 1f).** Tumors and metastases are depicted in **Fig S4ab.**

Finally, we tested the NAM+GEM treatment in the orthotopic KPC model. Briefly, 10^5^ KPC tumor cells were injected into the pancreas (the tumors in this model are more aggressive than in the orthotopic Panc-02 model because of the homozygous *p53^R172-/-^* mutation), and 10 days later the treatment with NAM+GEM was started (at this time point tumors are palpable in the pancreas) and continued for 14 days as outlined in **Fig S1c.** As shown in **Fig 1gh,** the tumor weight in the NAM+GEM group was significantly reduced (p<0.05) as well as the number of metastases (p<0.01), while the ascites production was high in all control groups (Grade 0-5), and minor in the NAM+GEM group (Grade 0-2) **(Fig 1i).** Tumors and metastases are depicted in **Fig S5ab.** Several mice died in the saline, NAM, or GEM group because of the aggressive cancer, but none in the NAM+GEM group.

To analyze the clinical relevance of the combination therapy, we tested the effect of NAM+GEM on the survival rate of orthotopic Panc-02 mice. Briefly, 10^6^ tumor cells were injected into the pancreas and 10 days later the NAM+GEM treatment was started and continued for 14 days as described above. Mice were monitored for the following weeks without any further treatment. Here we show that NAM+GEM significantly improved the survival time of the orthotopic Panc-02 mice not only compared to the saline group but also compared to all other control groups **(Fig 1j).**

### NAM+GEM increases the influx of immune cells and activates T cells while reducing TAM and MDSC

In skin cancer prevention, it has been shown that NAM significantly increased peritumoral and infiltrating CD4 and CD8 T cells compared to a placebo^23^. Here, we tested the effect of the combination of GEM+NAM on immune cells in the orthotopic Panc-02 tumors. At the end of treatment, whole tumors were digested with collagenase and dispase to generate single cell suspensions as we described previously^16^. Subsequently, the cell population was analyzed by flow cytometry. We found that the CD45^+^ cells (leucocytes) (p<0.0001) and CD8^+^ T cells (p<0.01) significantly increased in the NAM+GEM group compared to the saline group, while the increase in CD4 T cells was less robustly **(Fig 2a).** We also analyzed the number of TAMs in the Panc-02 tumors and found a significant reduction in the NAM+GEM group, but not in the GEM only group, compared to saline (p<0.05)**(Fig 2b).** The MDSC population was also reduced in tumors of the NAM+GEM group, although this was not statistically significant because of high variability in the saline treated mice **(Fig 2c)**. However, the MDSC population in blood was reduced by GEM alone (p<0.05)**(Fig 2d),** supporting our results in a previous study^16^.

**Figure 2:**
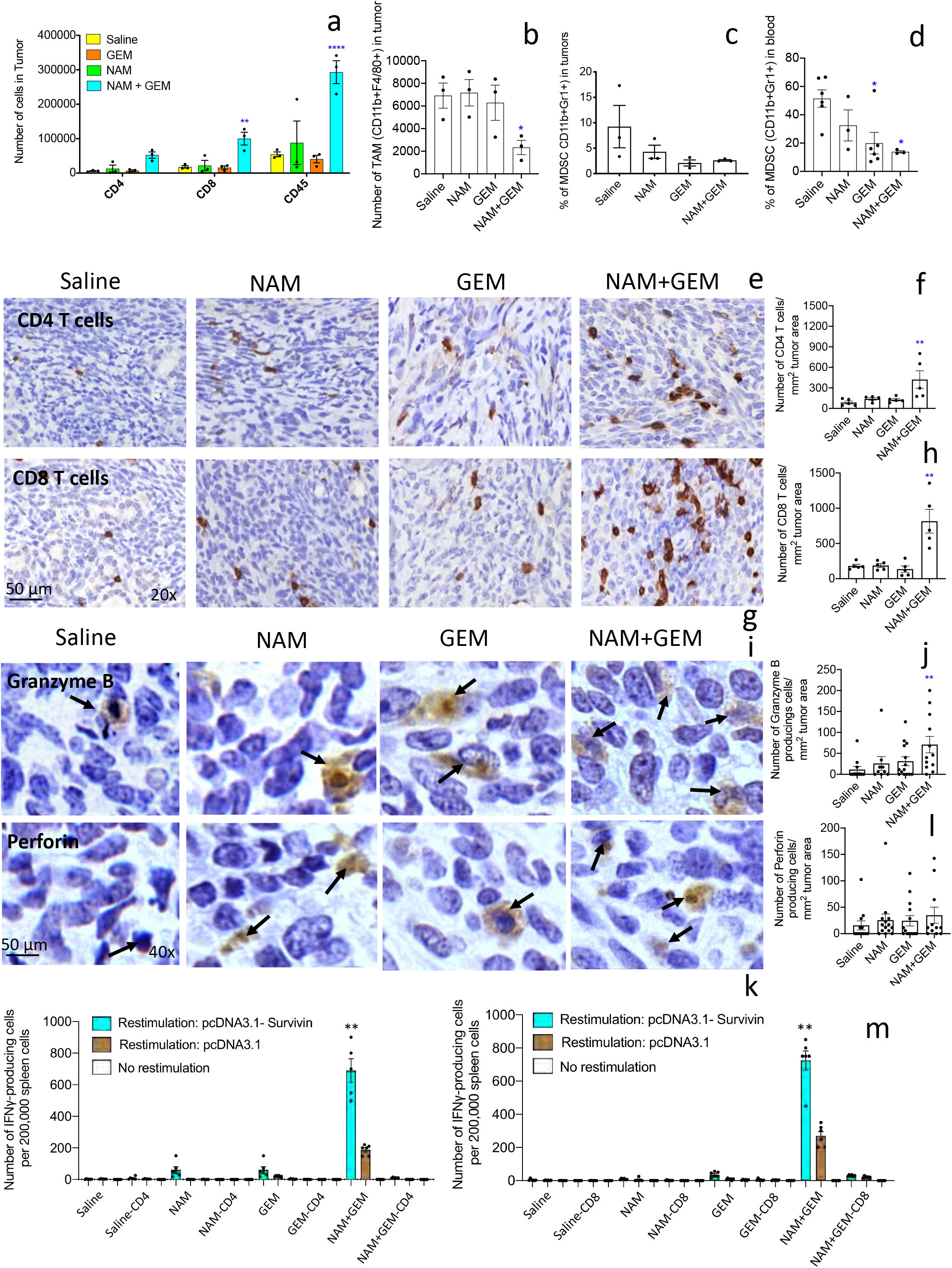
NAM+GEM increases the influx of immune cells and activates T cells while reducing TAM and MDSC. Immune cells (CD45+) were isolated from whole tumors and analyzed by flow cytometry for the presence of CD4 and CD8 T cells **(a),** TAM **(b)** and MDSC **(c).** In addition, blood was also analyzed for MDSC **(d).** Subsequently, tumor tissues were analyzed by IHC for the presence of CD4 and CD8 T cells **(e and g),** and quantified **(f and h).** 5 fields in each group were analyzed and the number of CD4 and CD8 T cells was calculated per mm^2^. n=5 mice per group. The results were averaged and analyzed by Mann-Whitney *p<0.05, **p<0.001 is significant. Granzyme B **(i)** and perforin **(k)** were analyzed by IHC, and quantified **(j and l).** 10 fields in each treatment group were analyzed and the number of perforin and granzyme B-producing cells was calculated per mm^2^. n=5 mice per group. The results were averaged and analyzed by Mann-Whitney *p<0.05, **p<0.001 is significant. Finally, spleen cells from NAM+GEM-treated and control mice were restimulated with survivin in an ELISPOT assay **(m).** CD4 and CD8 T cells were depleted by magnetic beads technology. The spleens of 5 mice were pooled in each treatment group. The spots of 6 wells were averaged and analyzed by Mann-Whitney **p<0.01 is significant.

To confirm the flow cytometry data we analyzed which T cells infiltrated the pancreatic tumors by immunohistochemistry, an approach we described previously^16^. As shown in **Fig 2ef and Fig S6ab**, the number of CD4 T cells was significantly higher in the tumors of orthotopic Panc-02 mice treated with NAM+GEM (p<0.01), but the number of CD8 T cells was more robustly increased compared to the saline group (p<0.01)**(Fig 2gh and Fig S6ab).**

We also analyzed the production of perforin and granzyme B in the pancreatic tumors through IHC. We found a significant increase in the production of granzyme B (p<0.01) **(Fig 2ij)** but less robust of perforin in the pancreatic tumors of mice treated with NAM+GEM compared to saline **(Fig 2kl+Fig S6c)**.

Since a strong decrease was observed in the size of the pancreatic tumors **(Fig 1de)** accompanied with a strong influx of CD4 and particularly CD8 T cells, we analyzed whether these T cells exhibited antitumor reactivity. For this purpose, spleen cells of NAM+GEM-treated and control mice were restimulated with survivin, a tumor-associated antigen (TAA) expressed by the Panc-02 tumors^33^, and analyzed by ELISPOT and magnetic bead technology. It appeared that CD4 and CD8 T cells were highly activated in the NAM+GEM group but not in the other groups **(Fig 2m).**

### NAM+GEM affects the stromal architecture with reduced extracellular matrix proteins and increased endothelial cells

The immunomodulatory effects observed in the above experiments demonstrates that T cell infiltration is improved in the NAM+GEM group. This compelled us to look at altered architecture of the tumor surrounding stroma such as the extracellular matrix. For instance, Collagen I, produced by CAFs, is known to prevent penetration of drugs and immune cells into the pancreatic tumors, particularly when cross-linked by hyaluronic acid fibrils. Here we showed a significant decrease (p<0.05) in Collagen I by Trichrome staining in the tumor area of NAM+GEM-treated mice compared to the saline group **(Fig 3abc and Fig S7a).** Moreover, we found a reduction in the expression of the hyaluronic binding protein (HABP) through RT-PCR in the NAM+GEM group compared to the saline group **(Fig 3d).** Also NAM or GEM separately reduced the expression of HABP.

**Figure 3:**
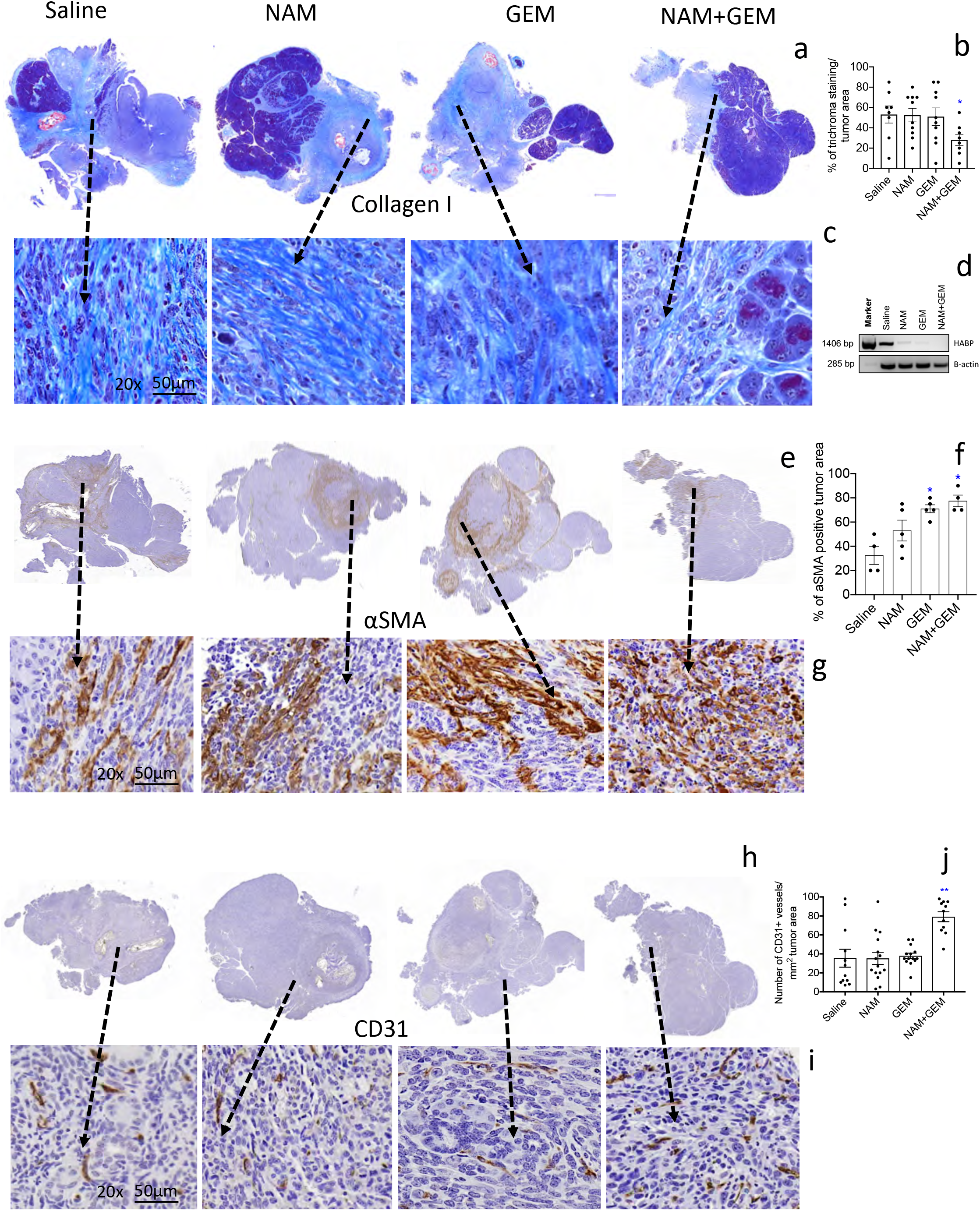
NAM+GEM decreases the production of Collagen I and HABP, and increases αSMA and CD31, in pancreatic tumors of orthopic Panc-02 model. Tumors were analyzed for Collagen I by trichrome staining **(a)** and quantified **(b),** and shown in more detail **(c).** n=5 mice per group and the results of 5 fields were averaged. Hyaluronic acid binding protein (HABP) fibrils was analyzed by RT-PCR **(d).** αSMA protein was by analyzed by IHC **(e),** and quantified **(f),** and shown in more detail **(g).** For both trichrome and αSMA the percentage of positive areas were determined and the results of n=5 mice per group was averaged. The presence of blood vessels in the pancreatic tumors was analyzed by IHC using anti-CD31 antibodies **(h)** and quantified **(j),** and shown in more detail **(i).** n=5 mice per group and the results of 10 fields were averaged. Mann-Whitney*p<0.05, **p<0.01 is significant. Error bars represent SEM.

αSMA, a protein expressed by a myofibroblastic CAF subpopulation, was significantly increased (p<0.05) in the tumor area of NAM+GEM-treated mice compared to the saline group **(Fig 3efg and Fig S7b).** Also, GEM-treated mice showed a significant increase in the αSMA compared to the saline group. Various reports have been published about the function of αSMA, which will be discussed later.

To allow the influx of immune cells into the tumors, blood vessels will be required. Therefore, we analyzed the tumors for CD31 expression. We found that CD31 was most abundantly expressed in the tumor areas of NAM+GEM treated mice **(Fig 3hij and Fig S7c).**

Peritumoral LNS are often observed in patients with PDAC. Some reported a correlation with improved outcome and others a correlation with a worse outcome^34,35^. Here we found peritumoral LNS in the NAM+GEM group (4/4) with CD4 and CD8 T cells **(Fig 4a),** and less frequently intratumoral LNS in NAM (3/5) and GEM (2/5) groups but not in the saline group **(Fig S8).** IHC showed CD31-positive vessels in both peritumoral and intratumoral LNS **(Fig 4b).** More detail about peri- and intratumoral LNS in the different treatment groups are shown in **Fig S8.**

**Figure 4:**
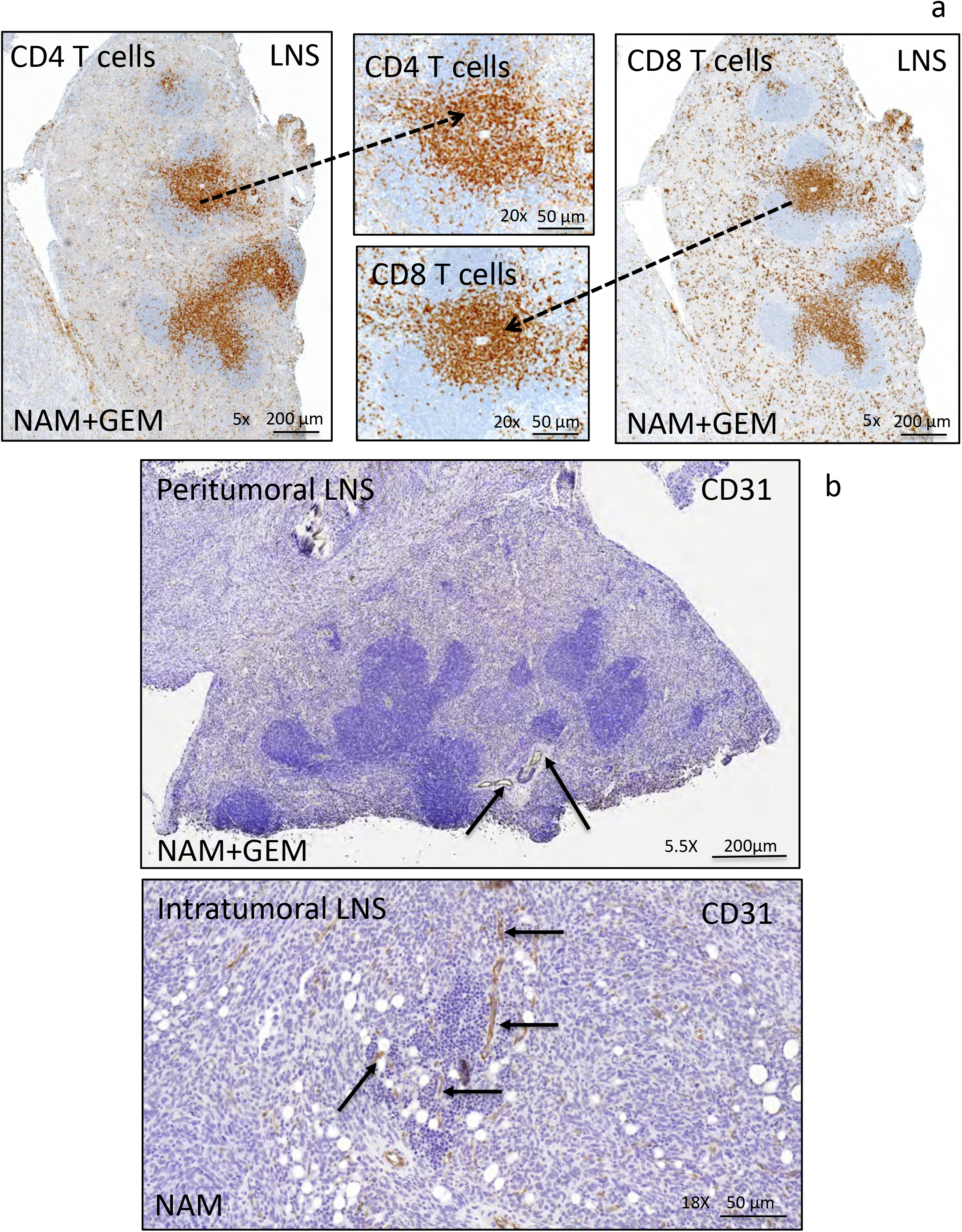
Peritumoral and intratumoral LNS in pancreatic tumors of orthotopic Panc-02 model. Detail of a peritumoral LNS **(a).** Peritumoral LNS, characterized by well-developed follicular structures with occasional germinal centers. CD4^+^ and CD8^+^ T cells are concentrated in the deep cortical zone/paracortex, where they were arranged in dense sheets, and are scattered within follicles and surrounding tissue. Detail of intratumoral LNS **(b).** Immunostaining for CD31 highlights the presence of vessels within peritumoral and intratumoral LNS in pancreatic tumors, which may allow the T cells to migrate to the tumor areas.

### Immuno-oncology gene expression profiling shows enhanced immune reaction and reduced pro-tumorigenic markers

To obtain more insight in the pathways potentially involved in the above described changes, we analyzed tumors of all treatment groups of the orthotopic Panc-02 model by Nanostring Technology and assessed specifically an immune-oncology gene panel **(Table S1).** We found significant changes in expression of genes relevant for immune responses and the progression/regression of the pancreatic cancer in the NAM+GEM group compared the saline group **(Fig 5a).** This includes, a 16-fold increase in the expression of Ccl21a, which is involved in T cell migration and recruitment of LNS, a 4-fold increase in IL1RL2, which is involved in activating T cells and increasing DC immunogenicity, a 10-fold increase in Fcer-1 which is involved in epitope spreading, a 4-fold increase in Ccl9 which is involved in attracting DC and activating T cells. Ccl21a was also increased in the NAM or GEM groups but less robust than in NAM+GEM. In another study, we also found that GEM improved the migration of T cells to pancreatic tumors^16^. Finally, we found a 4-fold reduction in Lcn-2 and a less robust but still significant reduction in Muc-1 and Epcam. These genes are involved in the promotion of invasive angiogenic drug resistant tumor cells, invasiveness and tumor progression (**Fig 5b).**

**Figure 5:**
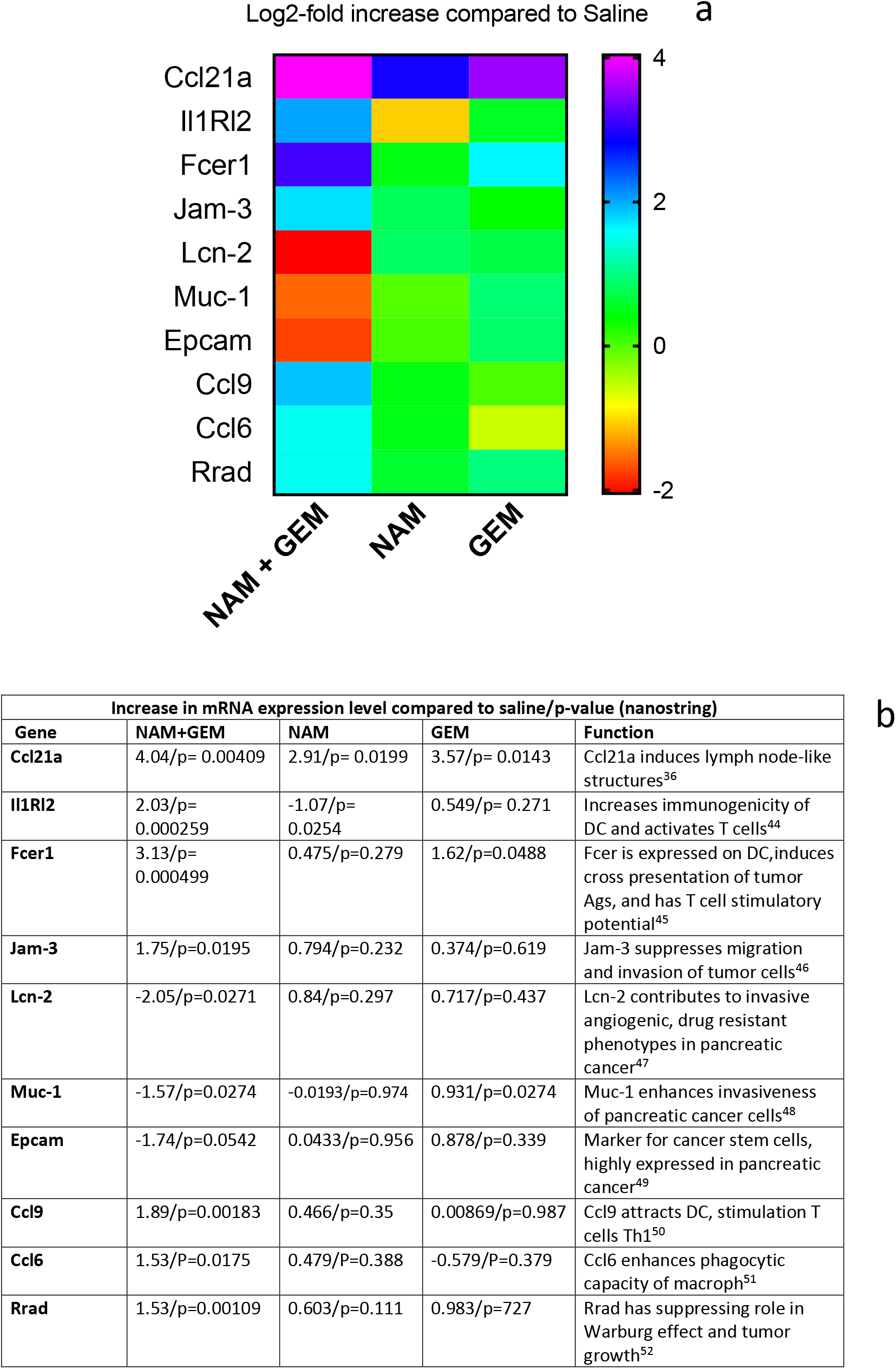
NAM+GEM increases the expression of genes involved in T cell migration and activation, and reduces the expression of genes involved in invasion of tumor cells. Gene expression profiles involved in T cell migration, activation, epitope spreading, invasion of tumor cells was analyzed by nanostring technology. A heatmap of relevant genes is shown **(a),** and the function of each gene and p-values **(b).** Gene expression levels in the NAM+GEM group were compared to the saline group and analyzed by ANOVA p<0.05 is significant.

## DISCUSSION

The success of cancer immunotherapies such as checkpoint inhibitors and CAR T cells in PDAC has been underwhelming, due to low immunogenicity of the tumor, strong immune suppression at multiple levels, and inefficient activation of T cells in the TME^8,9,11–14^. Here, we demonstrate that a novel combination of NAM+GEM in mice with pancreatic cancer not only reduced the pancreatic cancer (tumors and metastases) but also significantly improved the survival time compared to all control groups. We found suggestive evidence that tumor cells were killed by T cells through epitope spreading. This is based on the strong CD4 and CD8 T cell responses to survivin, which is expressed by the Panc-02 tumor cells^25^, and the 10-fold increase in Fcer in the tumors of NAM+GEM treated mice, which is responsible for epitope spreading. We also found that Ccl21a and IL1Rl2 were highly upregulated in the tumors of the NAM+GEM group compared to the saline group, which may be responsible for the T cell migration and activation and the formation of LNS^36,37^. A schematic view of potential mechanisms of NAM+GEM in pancreatic cancer is shown in **Fig 6.**

**Figure 6:**
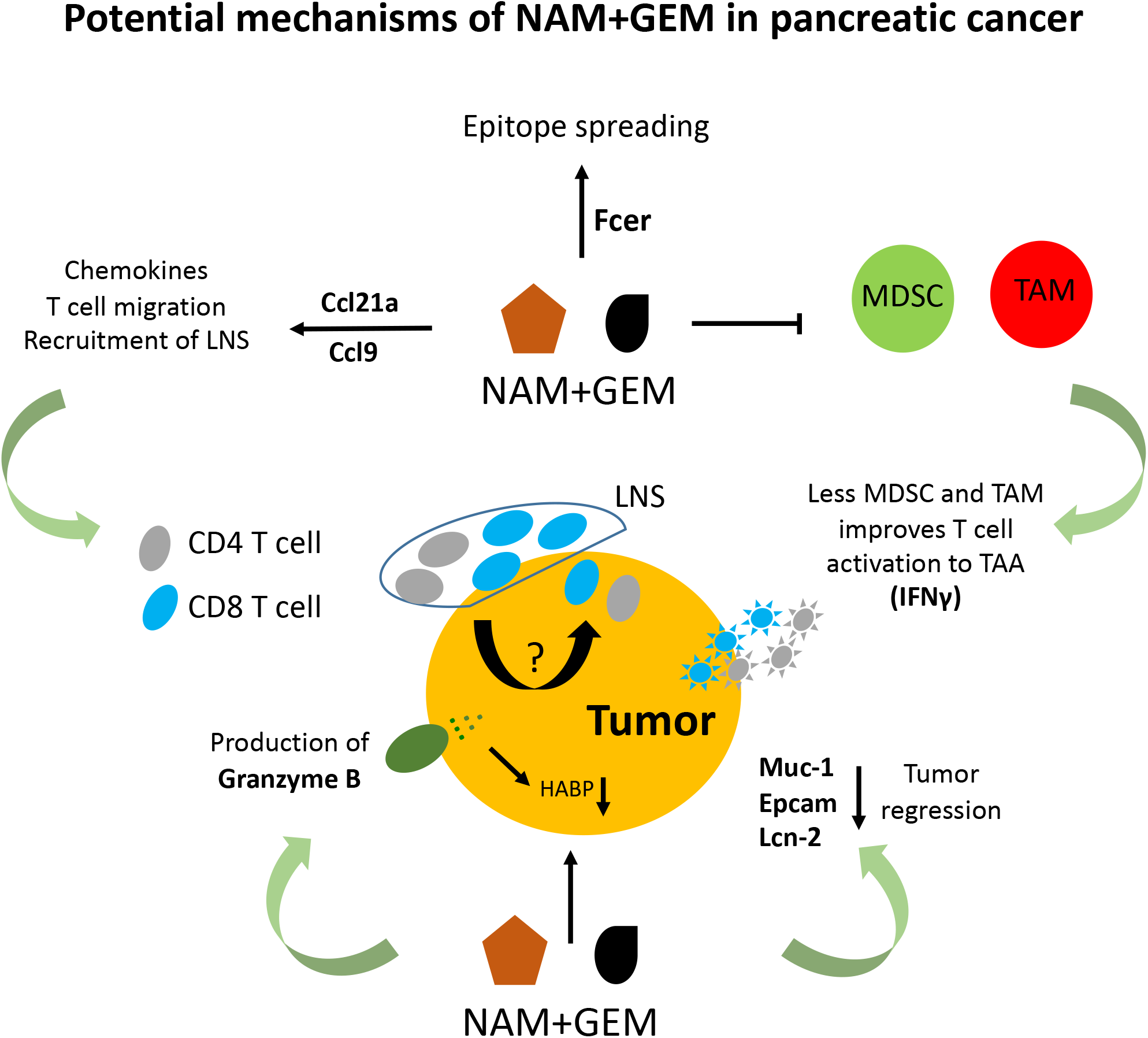
Potential mechanisms of NAM+GEM in pancreatic cancer. NAM+GEM increased the expression of Ccl21a and Ccl9 in the pancreatic tumors, which are involved in T cell migration and the formation of LNS through chemokines. We also found the production of Granzyme B in the tumors of NAM+GEM-treated mice, which may be responsible for decrease in the expression of HABP (fibrils linking the collagen I fibers), and the subsequent influx of T cells into the pancreatic tumors. NAM+GEM reduced the MDSC and TAM population in the pancreatic tumors, which resulted in improved T cell activation. NAM+GEM also increased the expression of Fcer which is involved in epitope spreading. Indeed, we found CD4 and CD8 T cells (producing IFN!) activated by survivin (expressed by Panc-02 tumors) through ELISPOT.

Immune suppression is one of the hallmarks of PDAC. Unfortunately, checkpoint inhibitors do not improve the outcome of therapy in PDAC patients, which we also observed when added to the NAM+GEM treatment. However, the TAM and MDSC populations significantly reduced in the pancreatic tumors or blood, respectively, of the NAM+GEM-treated mice compared to the saline group. This most likely has contributed to the improved T cell responses to survivin in the NAM+GEM group as well.

An interesting observation was the increased number of perforin and particularly granzyme B-producing cells in the tumors of the NAM+GEM group. While initially we assumed that this would be T cells, histopathological analysis (IHC) showed sparse large cells staining positive for perforin/granzyme B-producing cells, suggestive of macrophages. Other studies have shown that the production of granzyme B plays an important role in ECM remodeling^38^. Granzyme B cleaves fibronectin, which is involved in the formation of Collagen I fibrils^38^. Collagen I is produced by CAFs^39^. These fibrils form cross-links between the collagen fibers, which in turn may prevent drugs and immune cells from penetrating into the pancreatic tumor^1,40^. Cleaving fibronectin leads to degradation of Collagen I density, and may lead to a better infiltration of immune cells. We observed a significant decrease in Collagen I density and in hyaluronic acid binding protein in the NAM+GEM-treated group, in correlation with an influx of immune cells into the pancreatic tumors, and increase in CD31 positive blood vessels in the tumor areas.

In addition, we found peritumoral LNS in the NAM+GEM group and sometimes intratumoral LNS in the NAM or GEM group. Both peritumoral and intratumoral LNS, exhibited CD4 and CD8 T cells and CD31 positive vessels. The peritumoral LNS are more structured with well-developed follicular structures, while the intratumoral LNS showed an accumulation of CD4 and CD8 T cells in the tumor areas. Interestingly, in the NAM+GEM group LNS were predominantly peritumoral and correlated with improved survival. Whether T cells migrate from these LNS to the tumor cells and kill the tumor cells, need to be analyzed in more detail.

High αSMA expression is characteristic of myofibroblastic CAFs^39^. However, different correlations of αSMA expression with survival have been reported. One study in PDAC patients reported that high expression levels of αSMA in tumor stroma promoted invasion and cellular migration or was associated with a worse outcome^41^, while others found that high expression of αSMA is associated with improved outcome^42^. Also, deletion of αSMA^+^ fibroblasts in transgenic mice led to invasive undifferentiated tumors and reduced survival^43^. In our study we found that αSMA increased in the NAM+GEM group compared to the saline group and this correlated with significant improved survival of the orthotopic Panc-02 mice. This heterogeneity and plasticity of αSMA in the reported studies may have been the result of different types of therapy. Therefore, more detailed studies are required to obtain better insight in the role of αSMA in PDAC in relation to different therapies. It is yet unclear how these data, that are snapshots in time, need to be interpreted in a context of CAF heterogeneity and plasticity^5^.

In summary, we have demonstrated that NAM+GEM significantly reduced pancreatic cancer in different mouse tumor models, and significantly improved the survival time compared to all control groups. This correlated with a significant more T cells in the pancreatic tumors, as well as a significant increase in the production of granzyme B, epitope spreading of survivin and perhaps other TAA, and reduction in TAM and MDSC in the tumor microenvironment. The results of this study, and its promising effect in a phase 3 clinical trial to prevent skin cancer^23^, suggest that NAM as addition to GEM, which is the backbone of standard care for PDAC patients, could be a promising new lead for the treatment of pancreatic cancer.

## Author’s Contributions

### Conception and Design

BC. Selvanesan, L.Meheus, I. Rooman, C.Gravekamp.

### Development and Methodology

BC. Selvanesan, K. Meena, A. Back, O. Lara, I. Rooman, C. Gravekamp.

### Acquisition of data

BC. Selvanesan, K. Meena, O. Lara, A. Beck, l. Meheus

### Analysis and interpretation of data

BC. Selvanesan, K. Meena, A. Back, O. Lara, I. Rooman, C. Gravekamp.

### Writing, review, and/or revision of the manuscript

BC. Selvanesan, O. Lara, I. Rooman, C. Gravekamp.

### Study supervision

I. Rooman and C. Gravekamp

## Acknowledgements

We greatly thank Ms. Hong Zhang, Department of Pathology, and Dr. Vera DeMarais, Director of Light Microscopy and Image Analysis, Department of Structural and Cell Biology, Einstein for providing outstanding training and support regarding the IHC and image analysis. **Funding:** This work was supported by the Anticancer Fund, a private donation of Janet and Marty Spatz, NCI cancer center support P30CA013330 (Flow Cytometry Core, Pathology Core), and Dr. Rooman was supported by the FWO Odysseus Program (Research Foundation Flnaders).

## SUPPLEMENTARY INFORMATION

**Figure S1:**
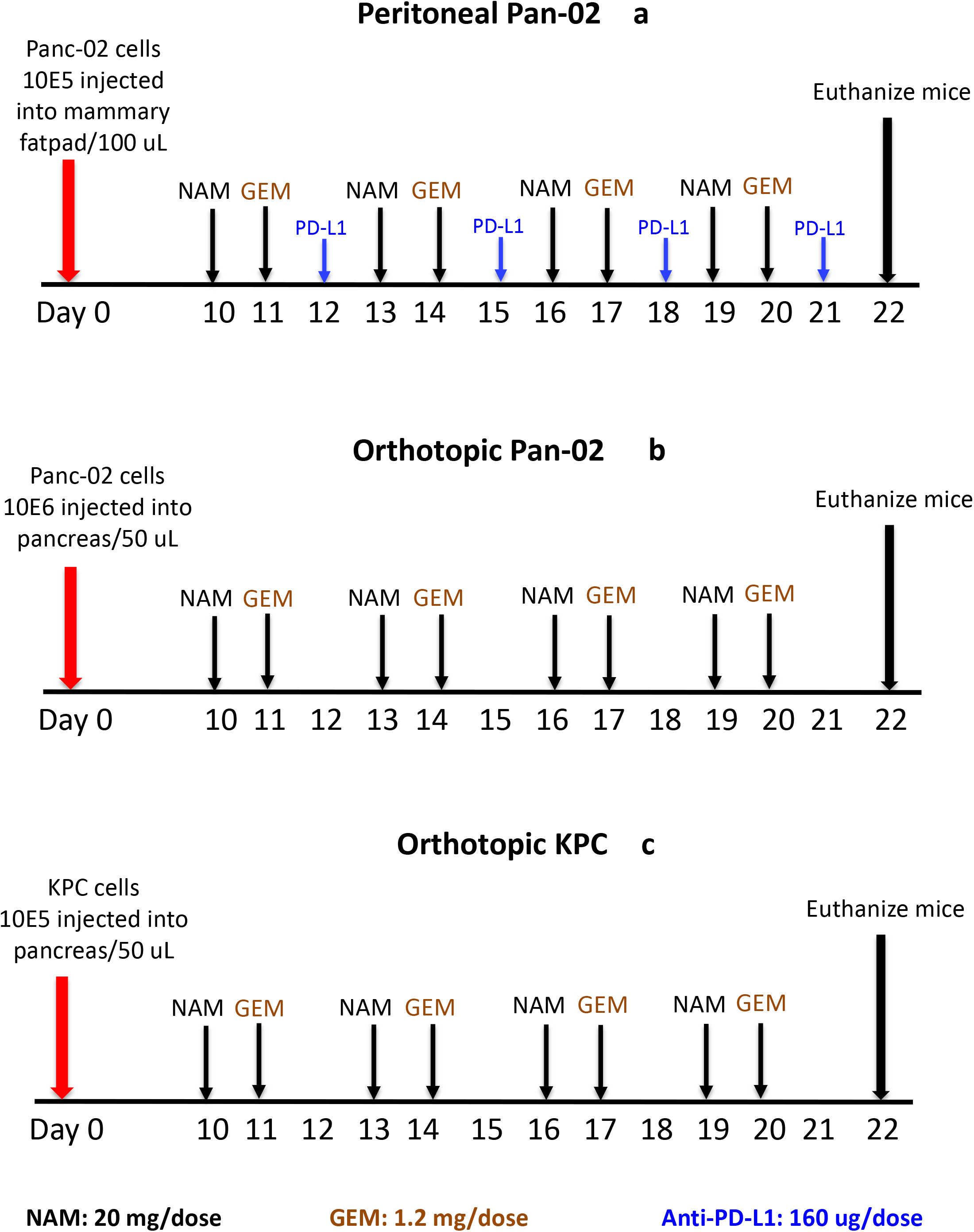
Generation of tumors and metastases and treatment protocol of NAM+GEM. Three different mouse models have been used. <u>Peritoneal Panc-02 tumor model **(a):**</u> 10^5^ Panc-02 tumor cells were injected into the mammary fatpad as we described earlier^1^. <u>Orthotopic Panc-02 model **(b):**</u> 10^6^ Panc-02 tumor cells were injected into the pancreas as we described earlier^2^. <u>Orthotopic KPC model **(c):**</u> 10^5^ tumor cells were orthotopically injected into the pancreas. Tumors and metastases are developed within 10 days after tumor cell injection. All treatments were started 10 days after tumor cell injection and continued for 2 weeks. NAM (20 mg/200 μL/dose) was administered orally every 3^rd^ day followed by GEM ip (1.2 mg/200 μL/dose) every 3^rd^ day and continued for 2 weeks. In one experiment anti-PDL-1 antibodies (160 μg/200 μL/dose was added to the combination therapy between each NAM and GEM dose.

**Figure S2:**
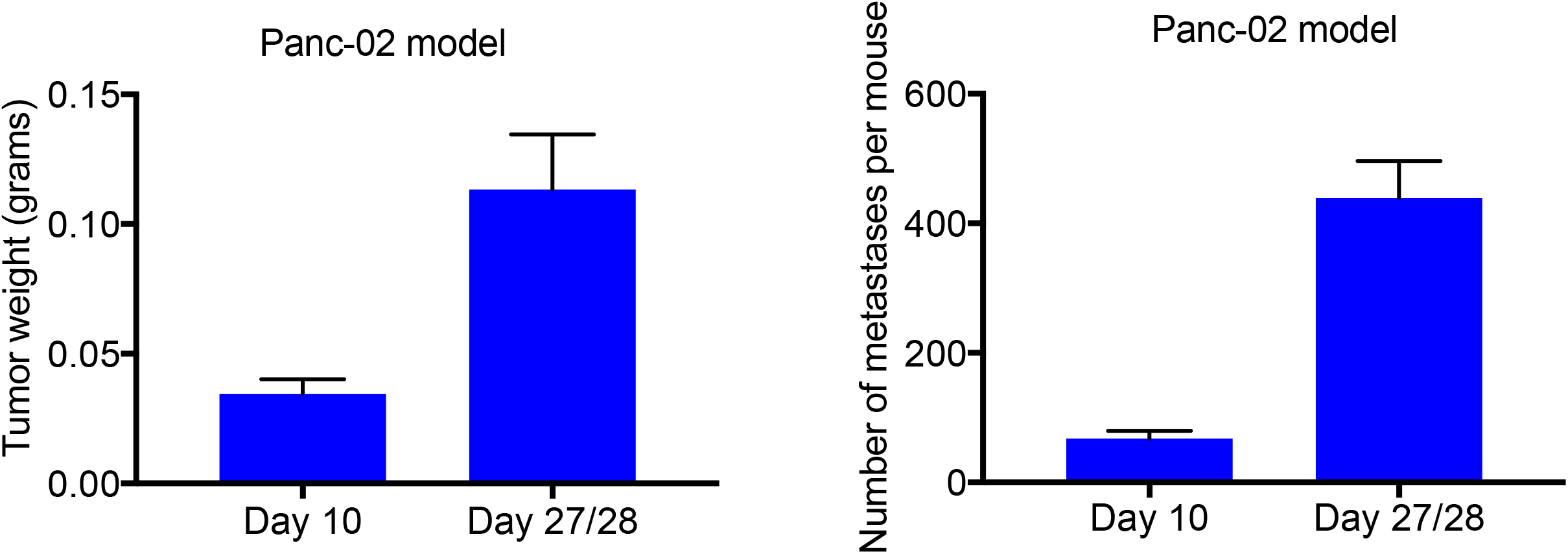
Tumor weight and number of metastases 10 days after tumor cell injection in peritoneal Panc-02 model. 10^5^ Panc-02 tumor cells were injected into the mammary fatpad as we described earlier^1^. Ten days later, we euthanized the mice and determined the tumor weight and number of metastases. The results of one experiment with n=5 mice per group was averaged. SEM=standard error of the mean.

**Figure S3:**
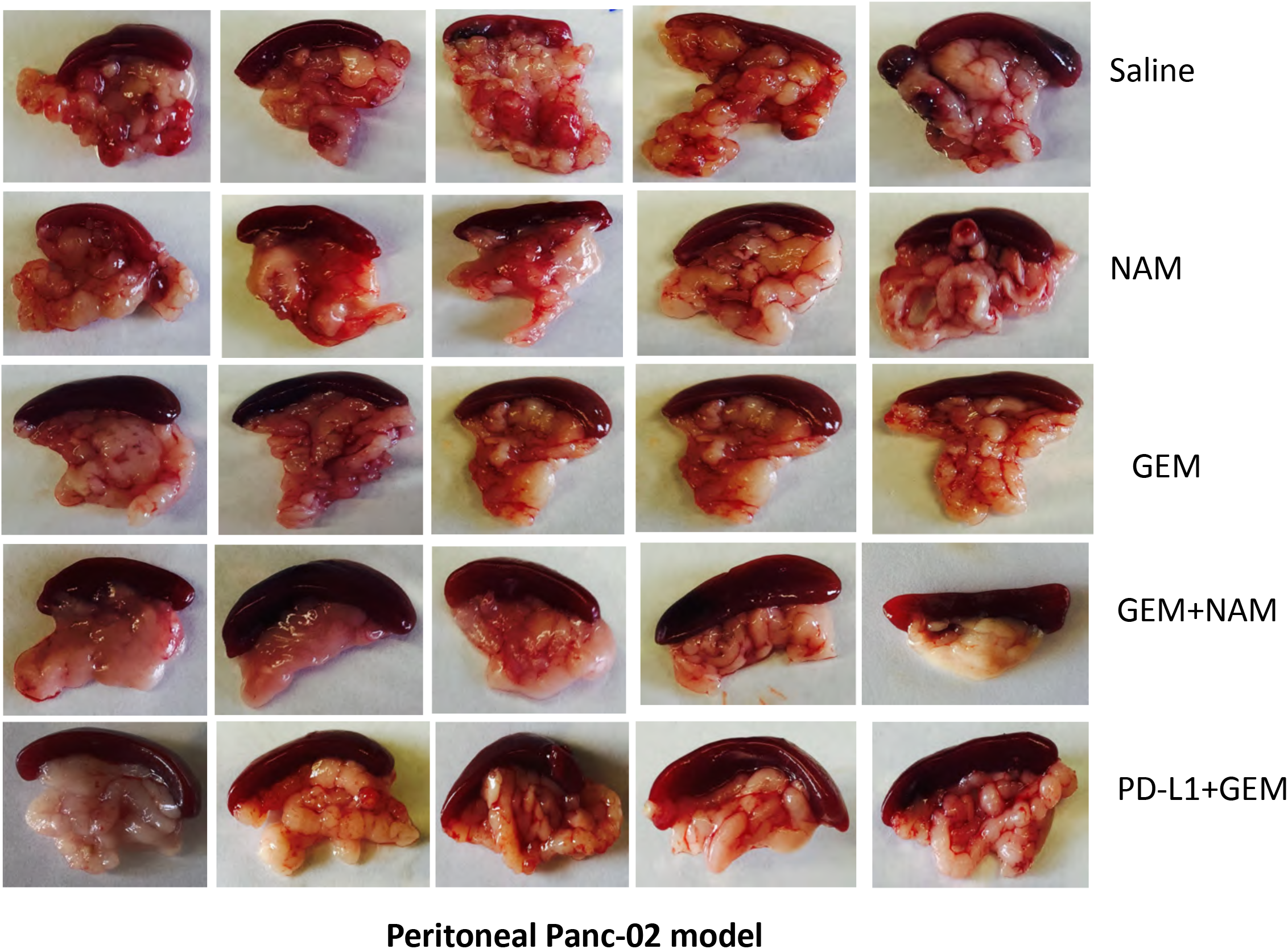
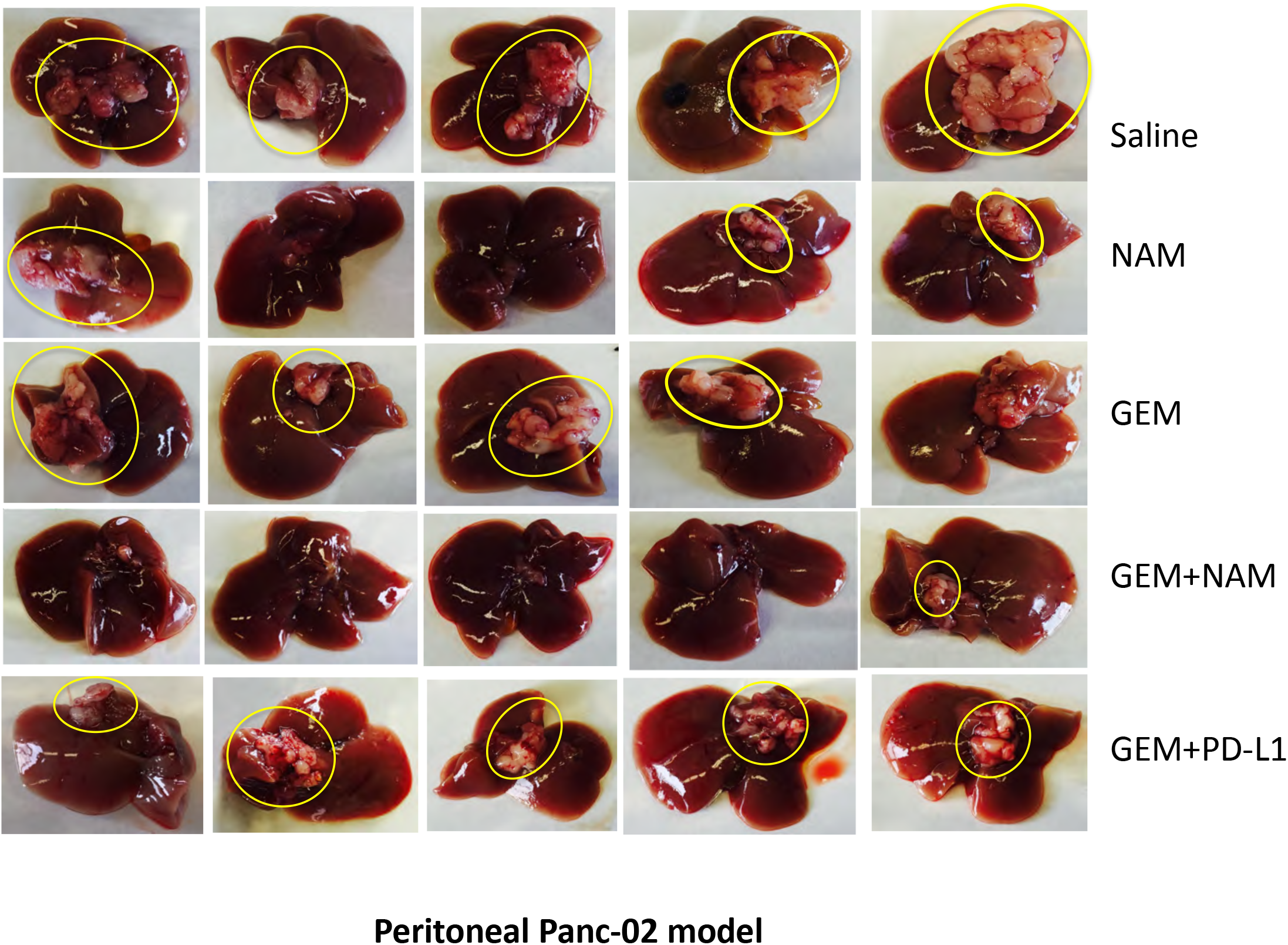
NAM+GEM reduce metastases in pancreas and liver in peritoneal Panc-02 model. Tumors and metastases were generated and treatments with NAM and GEM as described in **Fig S1a.** Here we show the metastases in the pancreas **(a)** and liver **(b)** of Panc-02 mice treated with NAM+GEM and control groups. In addition, anti-PD-L1 treatment was added to the NAM+GEM combination therapy.

**Figure S4:**
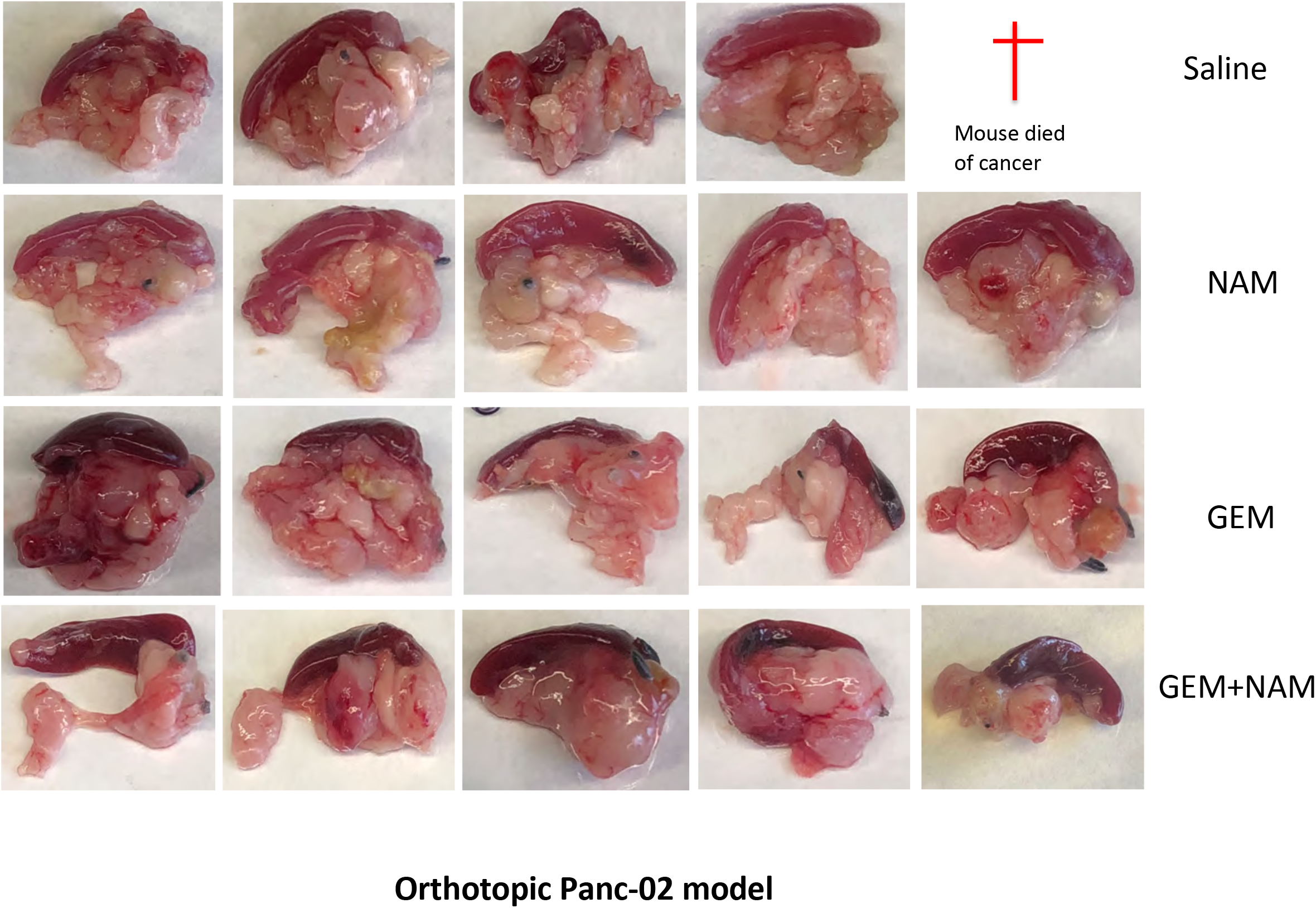
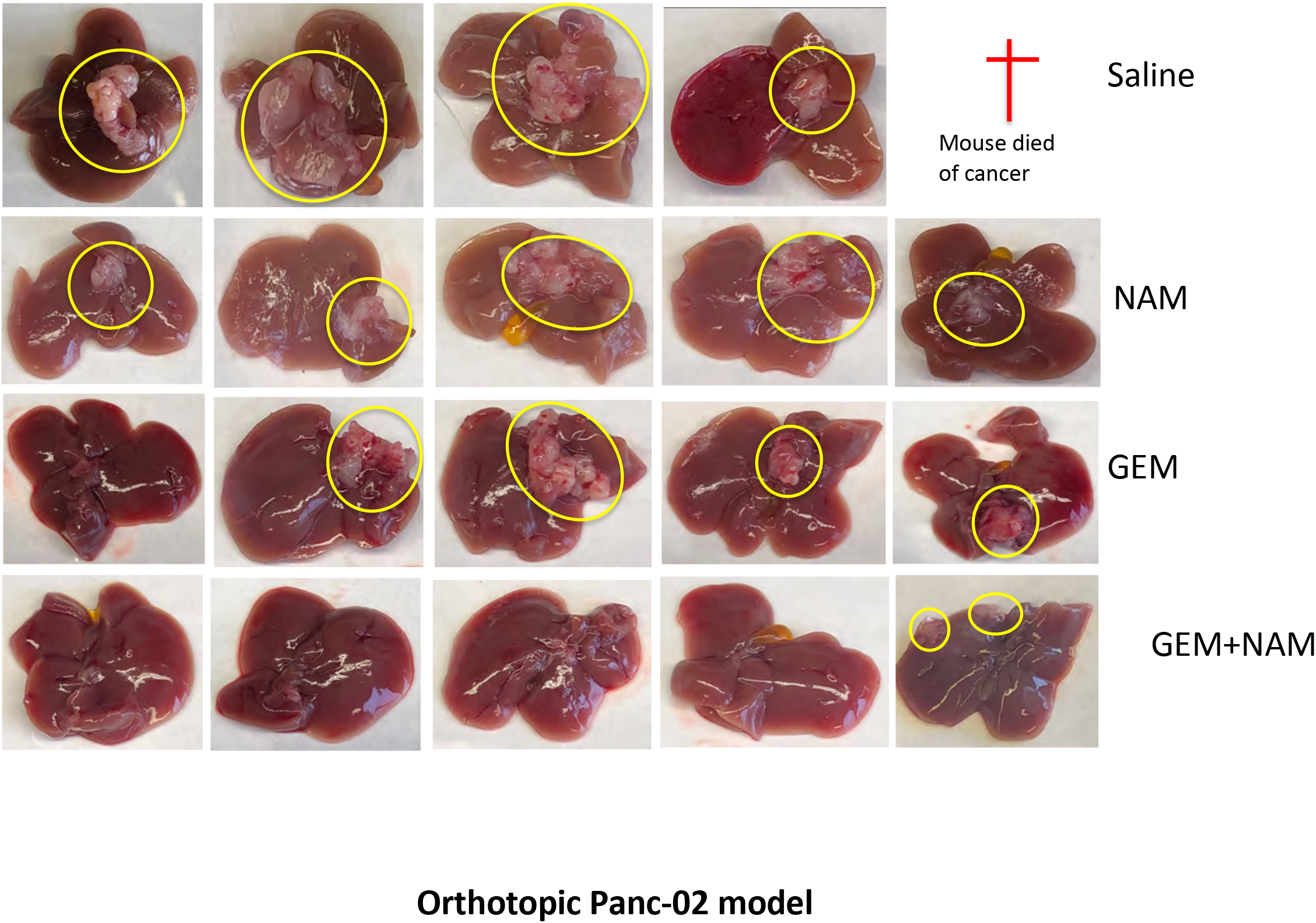
NAM+GEM reduce pancreatic tumors and metastases liver in orthotopic Panc-02 model. Tumors and metastases were generated and treatments with NAM and GEM as described in **Fig S1b.** Here we show tumors in the pancreas **(a)** and metastases in the liver **(b)** of Panc-02 mice treated with NAM+GEM and control groups.

**Figure S5:**
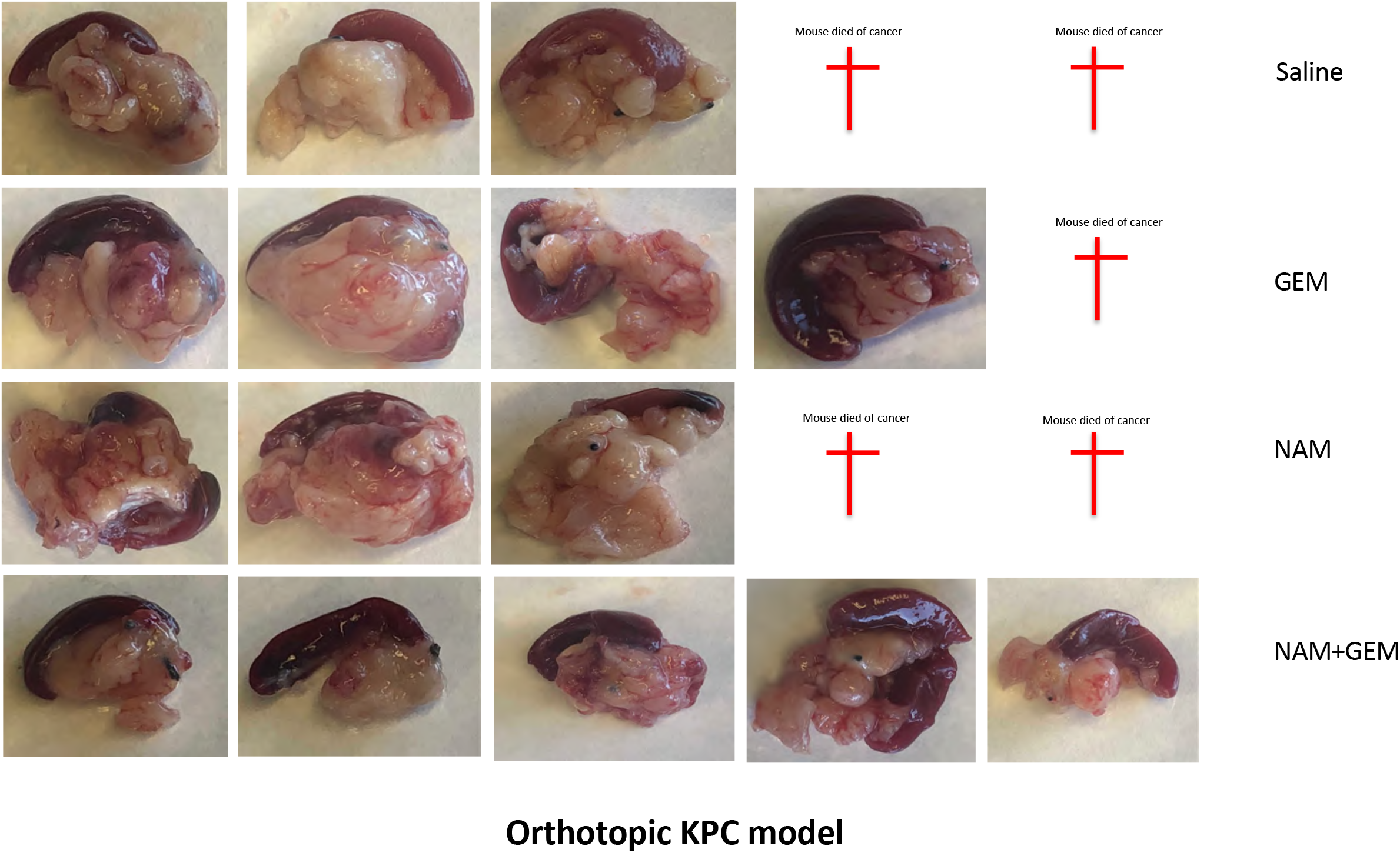
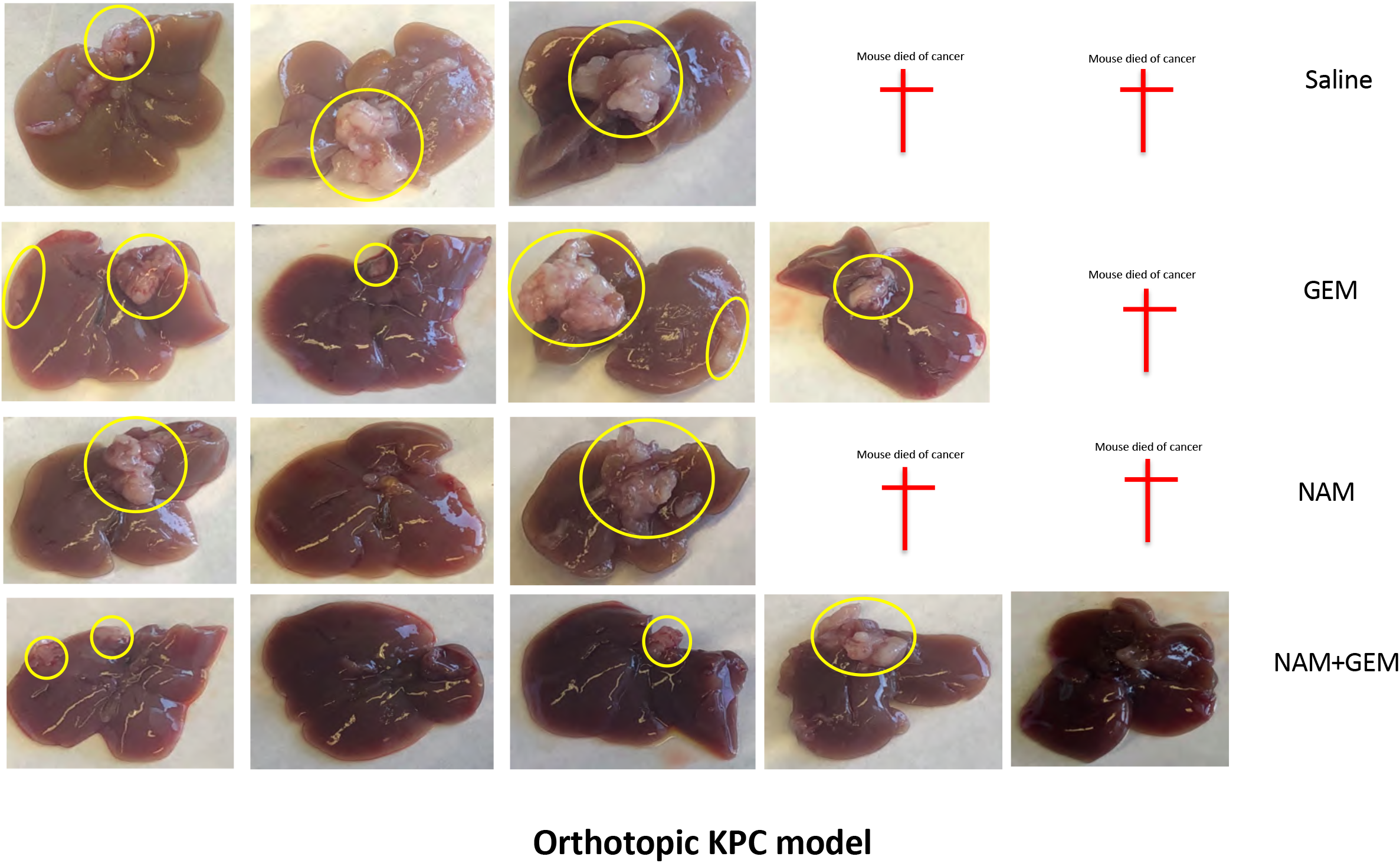
NAM+GEM reduce pancreatic tumors and metastases liver in orthotopic KPC model. Tumors and metastases were generated and treatments with NAM and GEM as described in **Fig S1c.** Here we show tumors in the pancreas **(a)** and metastases in the liver **(b)** of Panc-02 mice treated with NAM+GEM and control groups.

**Figure S6:**
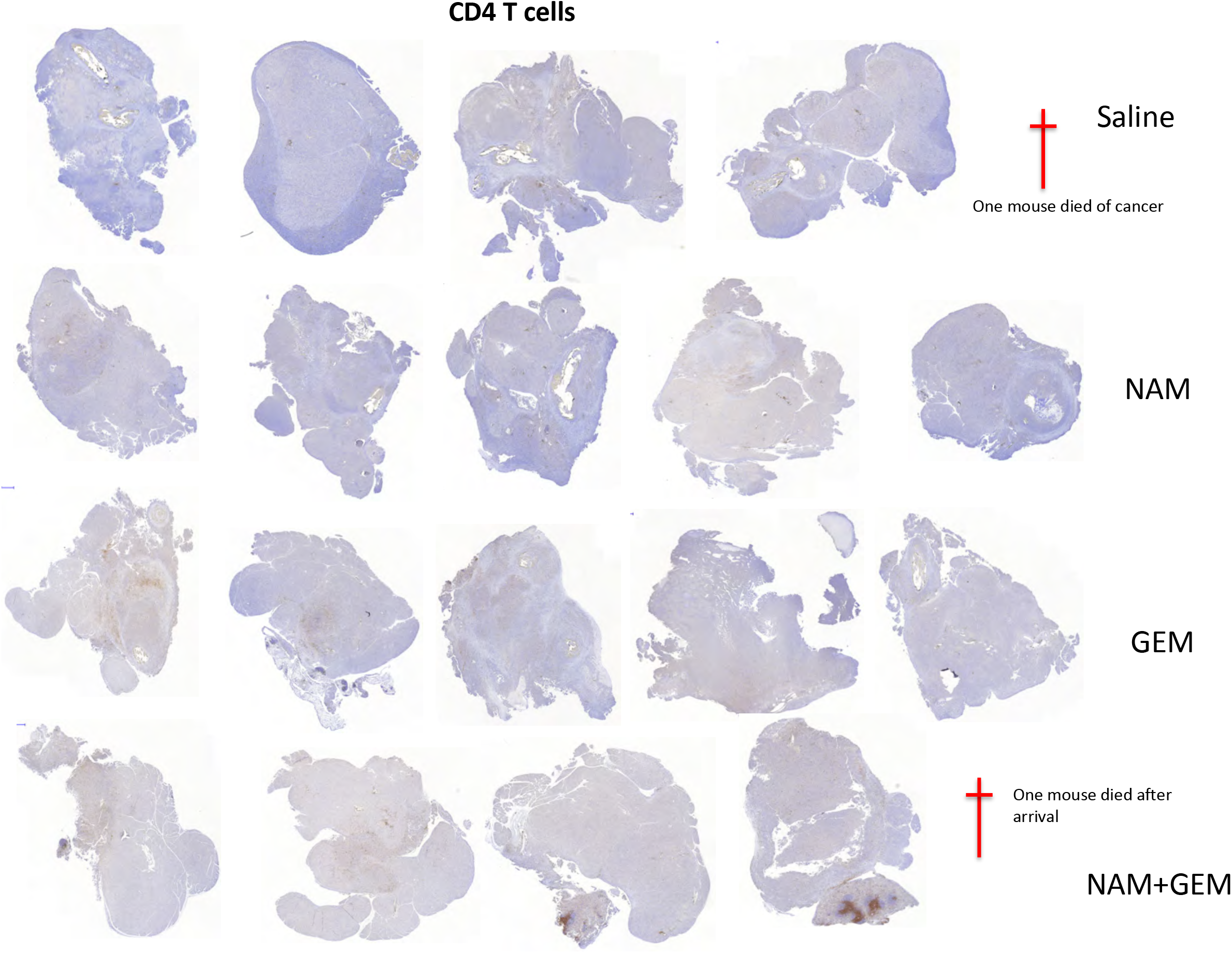
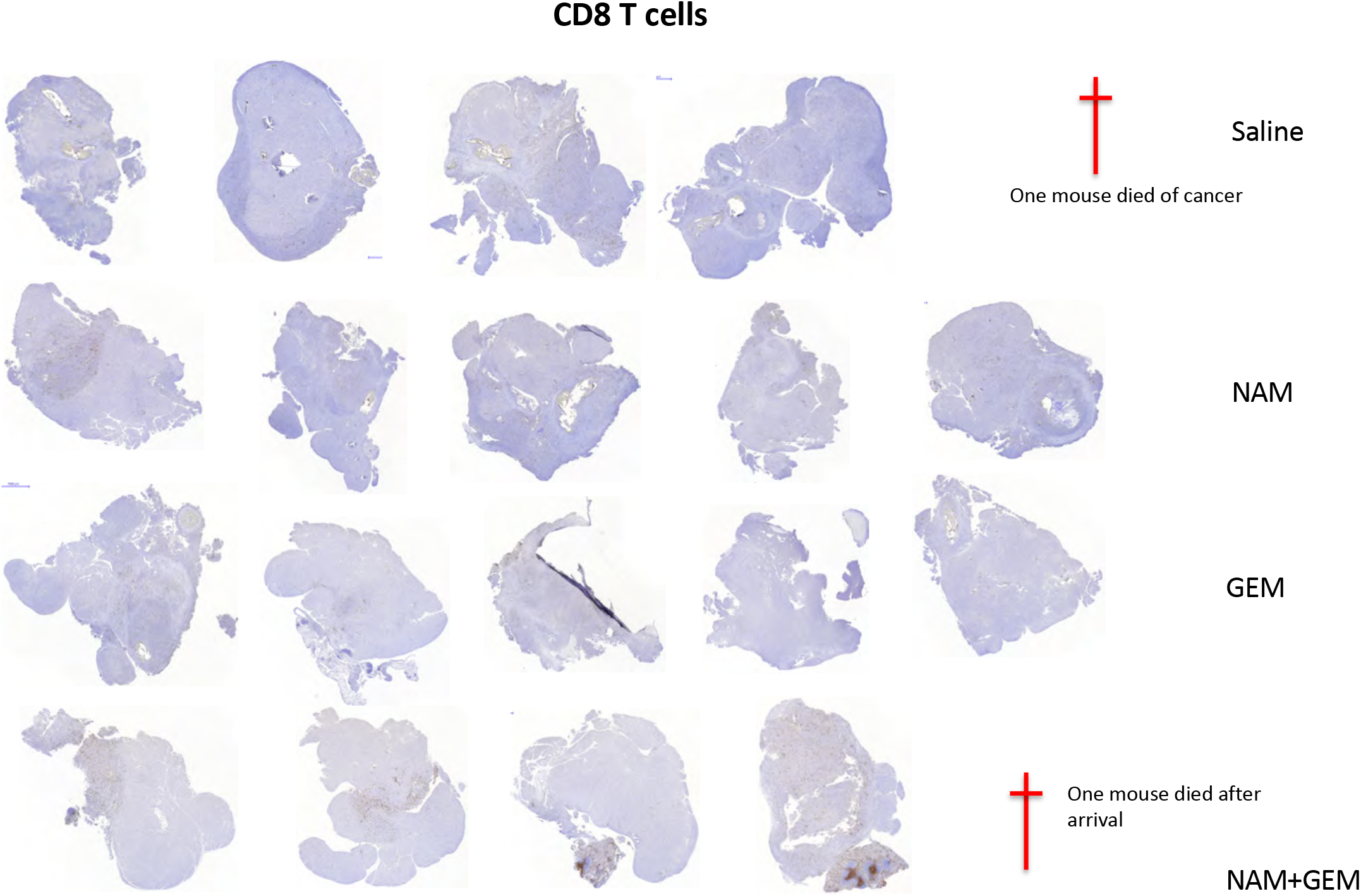
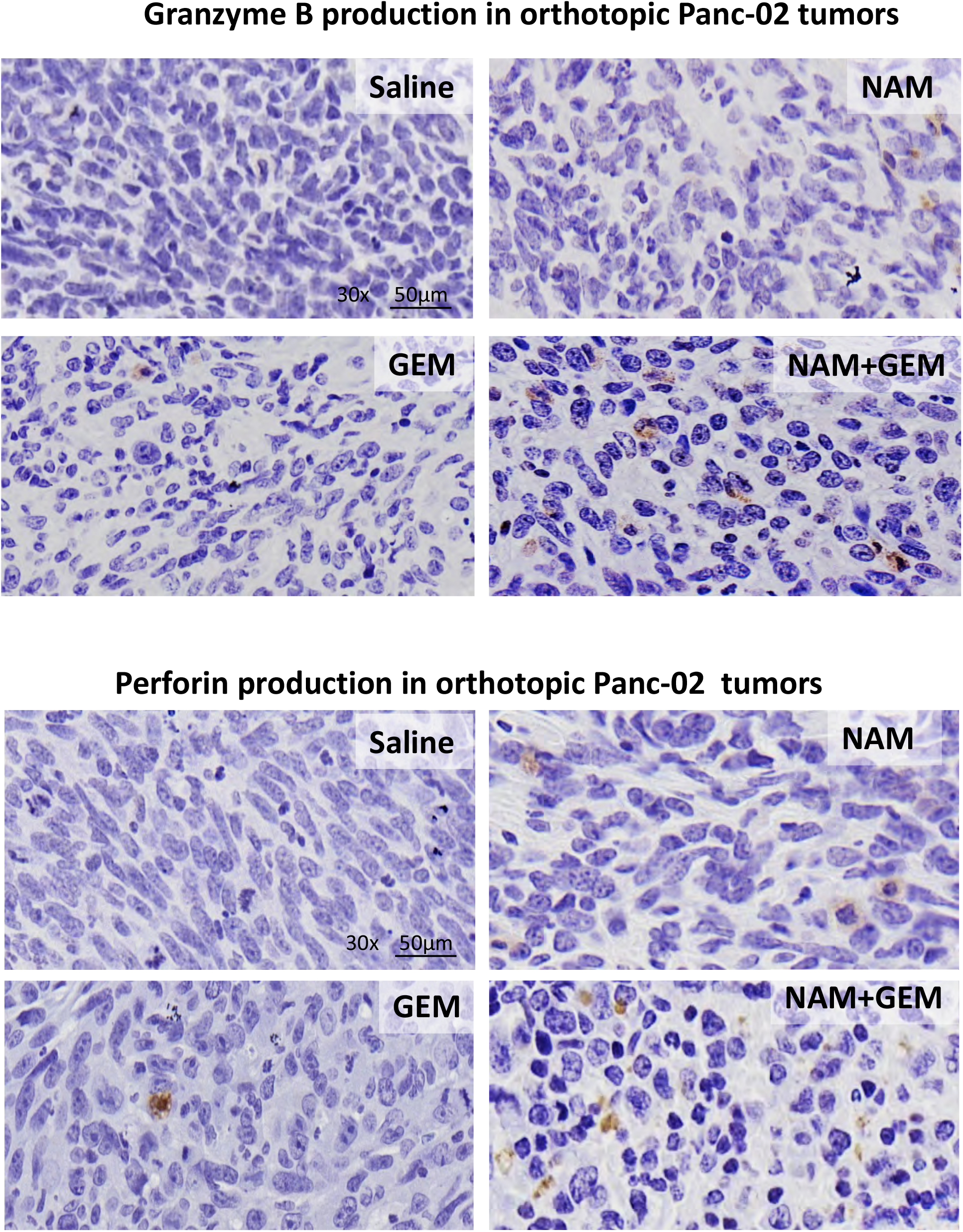
CD4 and CD8 staining of pancreatic tumors of orthotopic panc-02 model of all treatment groups. Overview of CD4 **(a)** and CD8 **(b)** staining by IHC. Stained sections are shown of each treatment group. Detail of perforin- and granzyme B-producing cells in pancreatic tumors analyzed by IHC **(c).**

**Figure S7:**
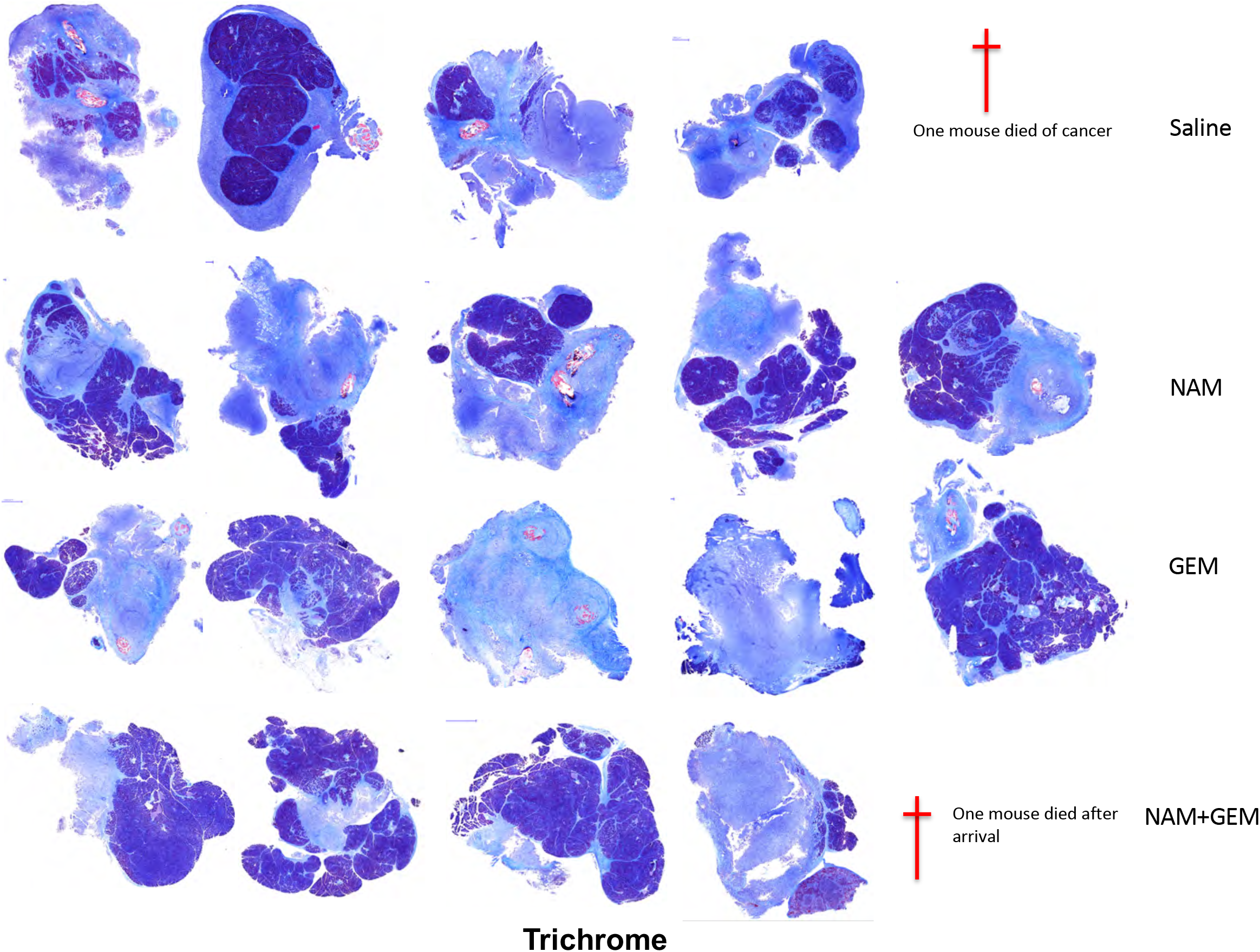
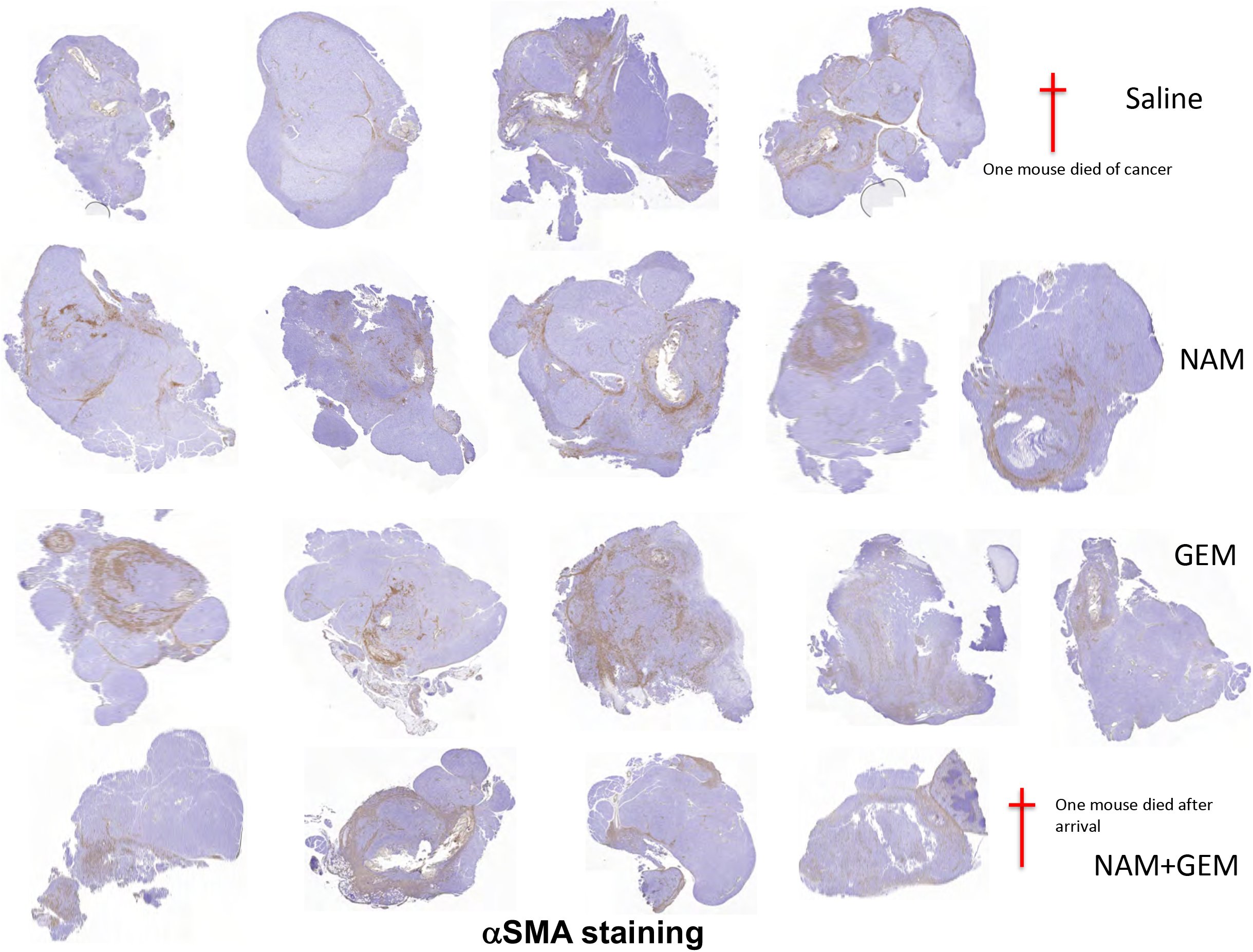
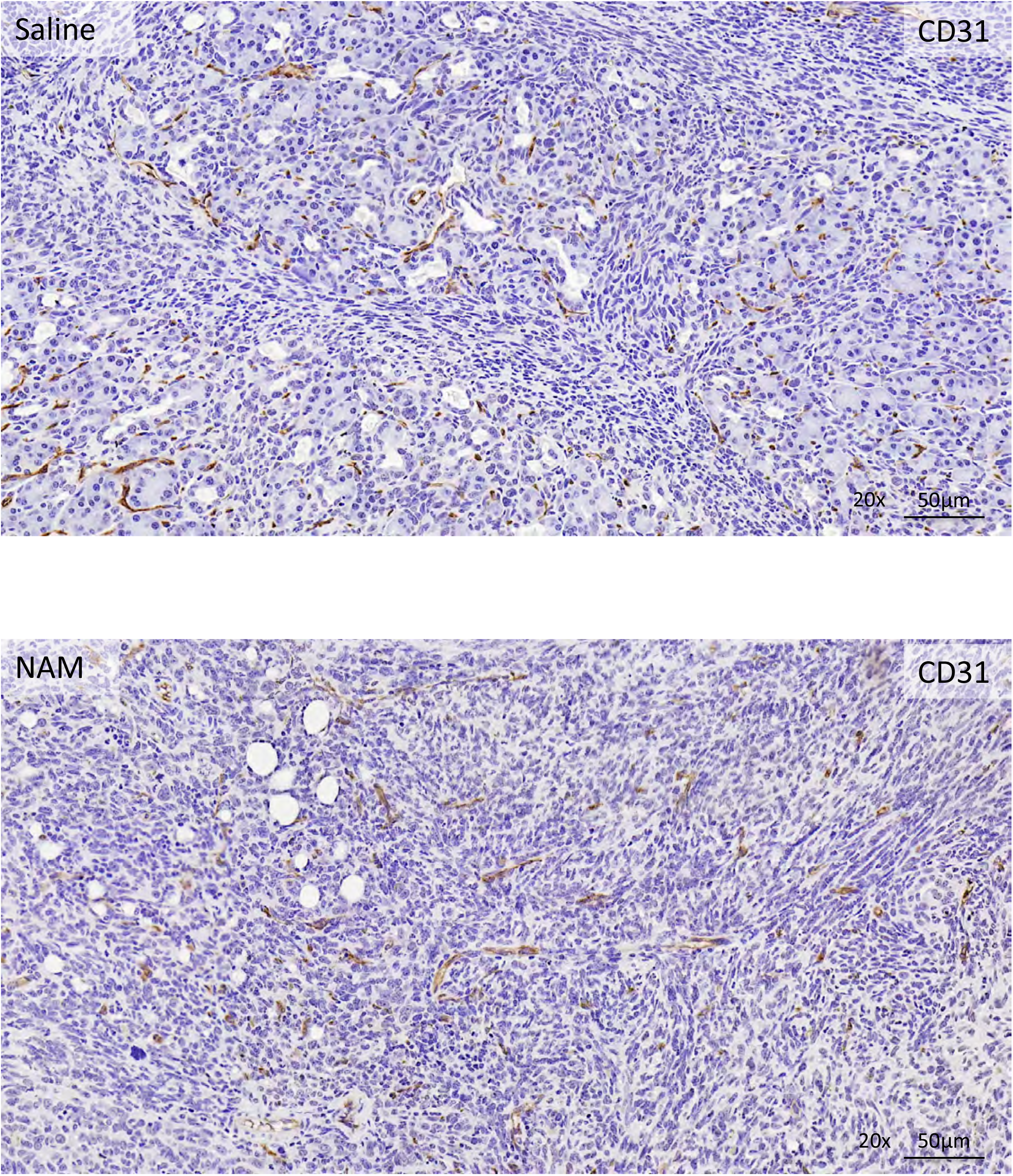
Trichrome and αSMA staining of pancreatic tumors of orthotopic panc-02 model of all treatment groups. **(a)** Overview of tumor sections of all treatment groups stained with trichrome. The dark blue areas represent the normal tissues (predominantly acinar cells), and the light blue areas the tumor tissues and tumor stroma. **(b)** Overview of tumor sections of all treatment groups stained with antibodies to α-smooth muscle actin (αSMA) protein. Detail of CD31 expression in pancreatic tumors by IHC **(c).**

**Figure S8:**
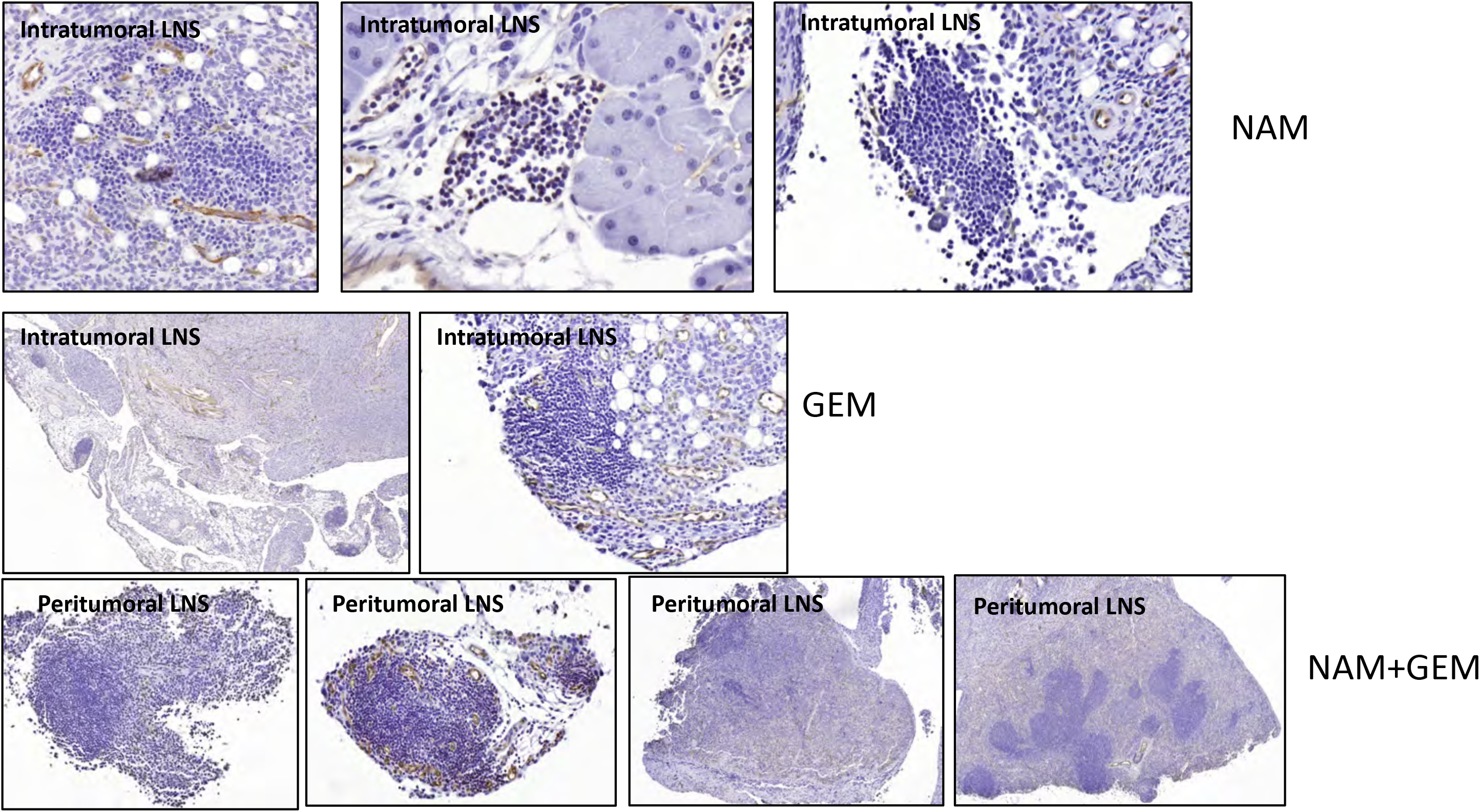
Peritumoral and intratumoral LNS in orthotopic Panc-02 model. NAM, GEM and NAM+GEM treatment groups exhibit formation of LNS, with peritumoral LNS most prominently developed in the NAM+GEM group. Both intratumoral and peritumoral LNS exhibit formation of vessels, as highlighted by CD31+ immunostaining, which may allow for migration of T cells to the tumors.

---

**Table S1:**
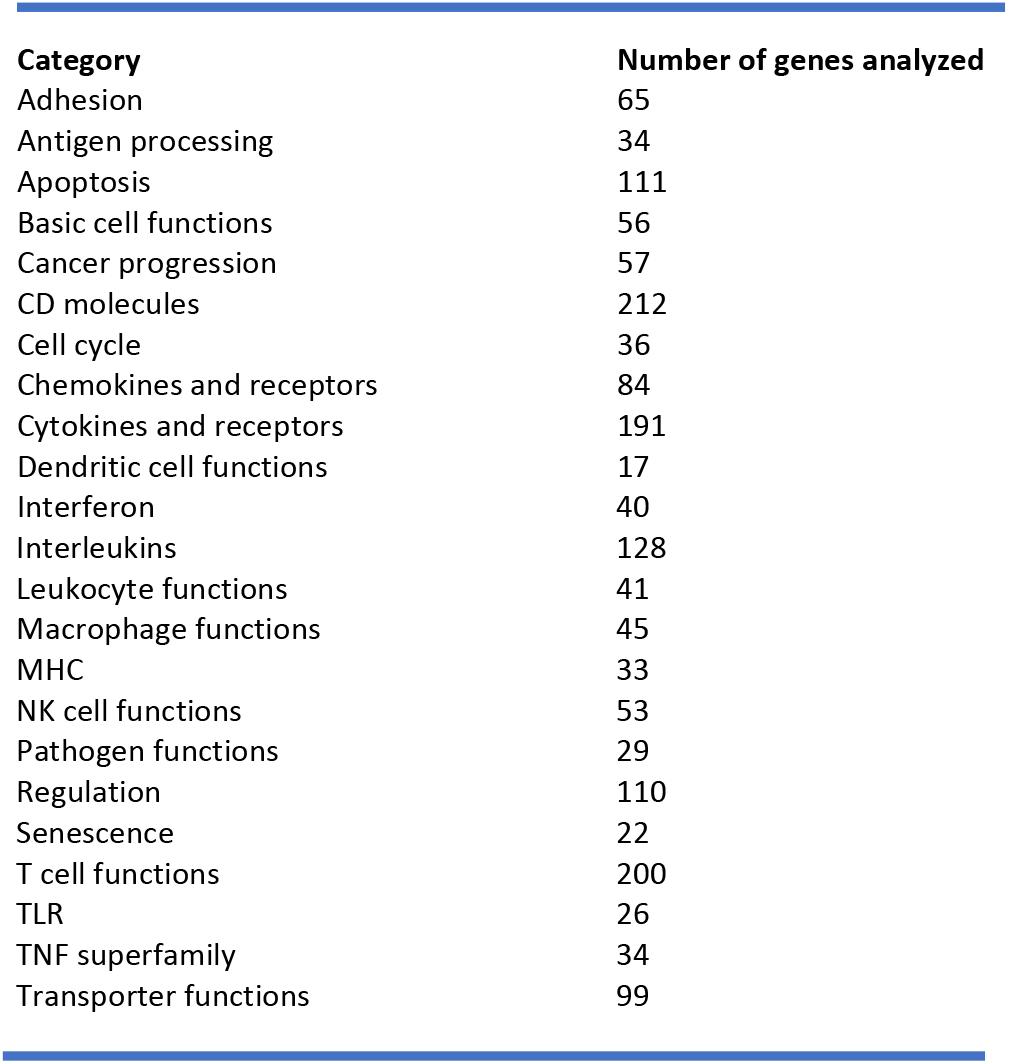
Pathways analyzed by Nanostring.

| Category | Number of genes analyzed |
| --- | --- |
| Adhesion | 65 |
| Antigen processing | 34 |
| Apoptosis | 111 |
| Basic cell functions | 56 |
| Cancer progression | 57 |
| CD molecules | 212 |
| Cell cycle | 36 |
| Chemokines and receptors | 84 |
| Cytokines and receptors | 191 |
| Dendritic cell functions | 17 |
| Interferon | 40 |
| Interleukins | 128 |
| Leukocyte functions | 41 |
| Macrophage functions | 45 |
| MHC | 33 |
| NK cell functions | 53 |
| Pathogen functions | 29 |
| Regulation | 110 |
| Senescence | 22 |
| T cell functions | 200 |
| TLR | 26 |
| TNF superfamily | 34 |
| Transporter functions | 99 |

